# Radiopaque Implantable Biomaterials for Nerve Repair

**DOI:** 10.1101/2023.01.05.522860

**Authors:** Kendell M Pawelec, Jeremy ML Hix, Erik M Shapiro

**Author notes:** Corresponding author: E. M. Shapiro.

## Abstract

Repairing peripheral nerve injuries remains a clinical challenge. To enhance nerve regeneration and functional recovery, the use of auxiliary implantable biomaterial conduits has become widespread. After implantation, there is currently no way to assess the location or function of polymeric biomedical devices, as they cannot be easily differentiated from surrounding tissue using clinical imaging modalities. Adding nanoparticle contrast agents into polymer matrices can introduce radiopacity and enable imaging using computed tomography (CT), but radiopacity must be balanced with changes in material properties that impact device function and biological response. In this study radiopacity was introduced to porous films of polycaprolactone (PCL) and poly(lactide-co-glycolide) (PLGA) 50:50 and 85:15 with 0-40wt% biocompatible tantalum oxide (TaO_x_) nanoparticles. To achieve radiopacity, at least 5wt% TaO_x_ was required, with ≥ 20wt% TaO_x_ leading to reduced mechanical properties and increased nano-scale surface roughness of films. As polymers used for peripheral nerve injury devices, films facilitated nerve regeneration in an in vitro co-culture model of glia (Schwann cells) and dorsal root ganglion neurons (DRG), measured by expression markers for myelination. The ability of radiopaque films to support nerve regeneration was determined by the properties of the polymer matrix, with a range of 5-20wt% TaO_x_ balancing both imaging functionality with biological response and proving that in situ monitoring of nerve repair devices is feasible.

## 1 Introduction

Peripheral nerve injuries (PNI) that result in the transection of nerve tissue, affect around 3% of trauma cases, such as those involved in motor vehicle accidents, and 30% of combat injuries, representing a significant healthcare burden on hospitals and patients [1,2]. With a gap between damaged nerve endings greater than a few millimeters, implants are required to guide regenerating nerve across the gap without applying a tensile force to the nerve stump, which is detrimental to nerve repair [3]. The devices clinically available for nerve repair are decellularized allografts or conduits made from either natural polymers, like collagen, or synthetic polymers, like poly(caprolactone) (PCL) and poly(lactide-co-glycolide) (PLGA) [4]. The benefit of synthetic polymer conduits is the lack of batch-to-batch variability inherent in naturally sourced materials, and their tunable degradation profile with a wide range of mechanical properties, depending on the polymer used. While these devices have demonstrated efficacy in the clinic [5], the use of biomedical devices carry inherent risks after implantation, such as movement or collapse over time [6]. Longitudinal non-invasive monitoring of devices may detect movement or collapse early and enable timely intervention to prevent catastrophic failure, but this is hampered by the fact that polymers in use for PNI repair cannot be radiographically distinguished from surrounding tissues by conventional clinical imaging modalities [7].

To enable longitudinal non-invasive radiological monitoring of polymeric PNI devices, contrast must be introduced into the polymer matrix. The use of nanoparticulate contrast agents incorporated into polymeric devices has been shown to significantly increase radiologists’ ability to non-invasively identify implant location and damage, using computed tomography (CT) imaging and magnetic resonance imaging (MRI) [8,9]. While both techniques are widely used in the clinic, CT is a readily available and cost-effective imaging modality, that can generate high-resolution 3D images with no limitation on tissue depth [10]. A number of nanoparticle contrast agents have been reported, based on metals or metal oxides. While extremely biocompatible nanoparticles from noble metals like gold or platinum are not cost-effective [10], other less expensive elements that generate high CT contrast include compounds and oxides of bismuth, barium and tantalum. Potential cytotoxicity concerns and fast excretion in native tissues have been raised for bismuth [11] and the highly water-soluble nature of barium sulfate (BaSO4), while commonly used as a clinical gastrointestinal contrast agent, presents hurdles in use with hydrophobic polymers. In contrast, tantalum oxide (TaO_x_) nanoparticles are biocompatible and inert [12], and can be differentially synthesized to be compatible with either aqueous media or hydrophobic systems [13].

While TaO_x_ nanoparticle contrast agents have been demonstrated to significantly improve radiopacity of synthetic polymers [8], little is known about their interaction with nerve tissue or during the complex process of nerve regeneration. Repair of PNI is highly coordinated, relying on glial cells to first bridge the injury and then guide regenerating neurons across the site. Finally, to reestablish functioning tissue, glia must remyelinate the neurites, ensuring efficient signal conduction [14]. During this process, glia are known to be sensitive to many environmental cues, including material chemistry, surface roughness and mechanics [15–17]. Addition of nanoparticles can potentially alter these properties, and thus, it is imperative to investigate how the contrast agent affects materials properties and simultaneously impacts the nerve repair process.

Three synthetic polymer matrices were chosen for study: PCL, PLGA 50:50 and PLGA 85:15, as these represent a broad range of mechanical properties and degradation profiles, and are already in use as clinical nerve repair technologies. Radiopacity was introduced to the polymers with the addition of 0-40wt% TaO_x_. The biological response of Schwann cells, the glia of the peripheral nervous system, was then investigated, examining attachment, proliferation and the ability to express markers indicating repair and remyelination. A range of TaO_x_ nanoparticle incorporation was identified that allowed both longitudinal monitoring via CT imaging and supported the nerve repair process. Functionalizing biomedical polymers with radiopaque contrast agents is a step towards the next generation of implantable devices that can accommodate non-invasive, long term radiological monitoring.

## 2 Materials & Methods

### 2.1 TaO_x_ nanoparticles

Throughout the study, TaO_x_ nanoparticles, with an aliphatic organosilane coating to impart hydrophobicity, were utilized. Manufacturing was based on our previously published literature [13], with the substitution of hexadecyltriethoxysilane (HDTES, Gelest Inc., cat no SIH5922.0) as the organosilane rather than (3-aminopropyl) trimethoxy silane (APTMS, Sigma).

### 2.2 Film manufacturing

Porous films were formed from polycaprolactone (PCL), poly(lactide-co-glycolide) (PLGA) 50:50 and PLGA 85:15. PCL (Sigma Aldrich) had a molecular weight average of 80 kDa. PLGA 50:50 (Lactel/Evonik B6010-4) and PLGA 85:15 (Expansorb^®^ DLG 85-7E, Merck) were both ester terminated and had a weight average molecular weight of 80-90 kDa. All three are biocompatible polymers used in clinically approved implants. Prior to casting films, polymers were solubilized in suspensions containing TaO_x_ nanoparticles in dichloromethane (DCM, Sigma); PCL was used at 4wt %, while PLGA solutions ranged from 10wt% (0-5wt% TaO_x_) to 8wt% (20-40wt% TaO_x_). To add porosity to the films, sucrose (mean particle size 31 ± 30 μm) was added to the solution, so that the final slurry was 70 vol % sucrose and 30 vol % polymer+nanoparticles. Suspensions were vortexed, then cast and dried on glass sheets. Dried films were removed and submerged in Milli-Q water to remove the sucrose, then dried for storage. Amount of TaO_x_ (0-40wt%) was calculated based on the weight percent of the total dry mass (polymer + nanoparticle). In addition, for cell culture, PCL films without porosity were created in the same way, without the addition of nanoparticles or sucrose.

### 2.3 Film characterization

TGA was used to characterize the weight percentage of TaO_x_ incorporated into polymeric scaffolds. Using a TA Q500 (TA Instruments), 12-14 mg of dry film was weighed in an alumina pan. The sample was heated from 20 - 600°C at a rate of 10°C/min under a nitrogen environment. The weight percentage of nanoparticles was calculated as the difference between the initial and final mass.

Film mechanics were characterized by tension tests on hydrated porous films. Samples (n=5) were cut from the films to be 5mm x 30 mm, and hydrated overnight in PBS at 37 °C prior to testing. To load the samples in the testing machine, the films were secured to card-stock frames, that were clamped into the grips, and then cut to allow sample movement. Tension tests were run at room temperature, at a rate of 1 mm/min until failure (TA XT plus texture analyzer). The elastic modulus was calculated as the linear portion of the stress strain curve. The yield stress and strain were found from a 0.2% off-set.

### 2.4 Glia cell culture and attachment

#### Preparation of films

Dry films were cut into 13mm diameter disks and were sterilized by immersing in 70% ethanol for 30 min, followed by two washes with sterile water. For in vitro cell tests, films were placed into 24-well inserts (CellCrown™, Z681903-12 EA). To reduce the leakage of media around the insert, each sample consisted of two layers: a bottom layer of non-porous PCL (cut to a circle of 24mm diameter) and a top layer of porous film (diameter: 13 mm). After assembly into the inserts, they were placed at the bottom of a 24 well tissue culture plate, and the entire assembly was sterilized for 30 min under UV light. Sterile inserts were stored at room temperature in a biological cabinet, in sterile water, until use. All coating was done under sterile conditions. Sterile films, pre-assembled into inserts, were coated with laminin (natural mouse laminin, Invitrogen (23017-015)). Each sample was incubated with 4 μg of laminin, diluted in sterile water, for 1 hr at room temperature. The surfaces were then washed and left in sterile water, at room temperature until used for cell culture, within 2 hr of coating.

Primary mouse Schwann cells were isolated from postnatal day 8 CD1 mouse nerves (ScienCell Research Labs, M1700), and used before passage 5 in all experiments. Glia were cultured in complete media (Dulbecco’s modified Eagle’s medium (DMEM, ThermoFisher Scientific, 11885), 10% fetal bovine serum (FBS, ThermoFisher Scientific, 160,000,036), 1% Penicillin– Streptomycin (ThermoFisher Scientific, 15,140,122), 21 μg/ml Bovine Pituitary Extract (BPE; Lonza, CC-4009), and 4 μM forskolin (Calbiochem, 344270)).

To quantify attachment onto films, 1×10^4^ cells/well were plated onto inserts in 400 μl complete media [15]. Samples were harvested 24 hr after seeding, by washing in phosphate buffered saline (PBS) and storing films at −80 °C for DNA quantification using Quant-iT™ PicoGreen™ dsDNA Assay Kit (ThermoFisher Scientific), per manufacturer’s instructions. Frozen inserts were digested in 200 μl papain buffer (0.1M phosphate buffer, 10mM L-cysteine, 2mM Ethylenediaminetetraacetic acid (EDTA), 3U/ml papain) at 60 °C overnight. In the assay, samples were diluted 1:4 in TNE buffer and loaded on a black 96 well plate. An equal amount of PicoGreen dye (1:200 in TNE buffer) was added to each well, and fluorescence was read at Ex: 480 nm, Em: 520 nm using a BioTek plate reader. Two standards were used: a cell standard (0– 5×10^4^ cells) of Schwann cells, created on the day of experimental cell seeding, and a DNA standard (1 μg/ml - 1 mg/ml) provided by the kit. All tests were done in triplicate, with two technical replicates each, and presented as mean ± standard deviation.

### 2.5 Co-culture model

The co-culture model simulating nerve regeneration was adapted from previously published literature [15].

#### Animal care

All animal handling and surgical procedures were performed in compliance with local and national animal care guidelines and approved by Michigan State University’s IACUC. For all experiments, 4 week-old CD1 female mice were used (Charles River Laboratories). Mice were group-housed in a 12-h light-dark cycle and provided with enrichment environment (EE) (e.g. nestlets, plastic huts); rodent chow (Teklad Global Diet^®^ 2918) and water (facility tap) was offered ad libitum.

#### Preparation of dorsal root ganglion (DRGs) cultures

Unless specified, naive DRGs at all rostro-caudal levels along the spinal cord were harvested immediately following euthanasia via CO2 inhalation. For each biological replicate, DRGs from 3-4 individuals were combined. DRGs were harvested, placed in ice-cold Leibovitz’s L-15 medium (Gibco), and digested in Collagenase type II (4 mg/ml, Worthington) and Dispase II (2 mg/ml, Sigma), diluted in L15 at 37 °C, for 45 min. Tubes were inverted every 10 min during enzymatic digestion. The digestion medium was pulled off and cells were rinsed with neural growth medium (DMEM/F-12/ GlutaMAX + 10% FBS + B27 supplement + Penicillin/Streptomycin).

#### Co-culture model

Porous films (0-40wt% TaO_x_) and glass coverslips (control surface) were coated with laminin, as before. Co-cultures consisted of 1×10^4^ Schwann cells per well, followed by the addition of 6-8×10^3^ naive DRG neurons per well, 4–6 h after initial Schwann cell seeding. During the first 24 h, the inserts were cultured in complete DRG medium. Following this, cells were cultured for 3 days in Schwann growth medium (DRG complete medium, 21 μg/mL BPE, 4 μM forskolin). Finally, the culture medium was replaced with differentiation medium (DRG complete medium, 50 mg/ml ascorbic acid, 21 μg/ mL BPE, 0.5 μM forskolin). Media was aspirated and replaced three times per week for 1 week. Samples were either fixed in 4% para-formaldehyde for 15 min and stored in PBS at 4 °C, for immunohistochemical staining (IHC), or inserts were placed in 200 μl Ripa buffer (Fisher Scientific) with 1× protease and phosphatase inhibitor cocktail (Sigma, PPC1010), and rocked gently overnight at 4 °C; the inserts were then removed and the protein lysis was stored at −20 °C until use.

### 2.6 Protein Expression

Protein was harvested from co-cultures in Ripa buffer and concentration was measured (BioRad, BCA kit). Expression of myelination markers was quantified using sodium dodecyl sulfate polyacrylamide gel electrophoresis (SDS-PAGE) and Western blotting. Full protocol details and images of complete blots are presented in Supplemental. Briefly, samples with 2× Laemmli buffer were incubated at 60°C for 20 min, and separated by SDS/ PAGE. After transfer to PVDF membranes (iBlot 2 Gel Transfer Device, invitrogen), membranes were blocked with 5% dried milk in PBST (phosphate buffered saline pH 7.4, containing 0.1% Tween-20) and probed with primary antibodies specific for Oct-6 (1:1,000, Aviva Systems Bio, ARP33061_T100), MPZ (1:1000; Aves, PZO), c-Jun (1:1000, Cell Signaling Technology, 60A8), Sox10 (1:1000, Abcam, ab155279), integrin α6 (abcam, ab181551), and integrin β1 (abcam, ab179471) at 4 °C overnight. The next day, the membrane was washed and incubated with anti-chicken IgY-HRP (12-341, Millipore) or anti-rabbit IgG-HRP (ab205718, abcam) at room temperature for 2 hr. Protein bands were visualized with Amersham ECL Plus substrate (cytiva, RPN2232) on a Li-Cor Odyssey FC and resulting signal was quantified via Image J.

Developed membranes were washed in PBS and stripped for 15 minutes in Restore Western Blot Stripping Buffer (Thermo Scientific 21059) before being either blocked and reprobed, or stained for total protein using the BLOT-Fast Stain (G Biosciences, cat# 786-34) according to manufacturer’s directions. The total protein signal was imaged under visible light on a c300 imager (azure biosystems). Protein expression was quantified via Image J and normalized to total protein to adjust the measured band intensity. For comparisons between membranes, all signals were normalized to PCL + 0wt% TaO_x_, which was run on every membrane. All data reported are the result of three independent replicates (two technical replicates each) and are reported as mean ± standard deviation.

### 2.7 Microscopy

Prior to staining, fixed samples were stored in PBS at 4 °C. Films were washed in PBS+0.1% Tween-20 (PBST), then permeabilized in 0.1% Triton X-100 in PBS, followed by two PBST washes. Samples were then blocked with 5 wt% bovine serum albumin (BSA) in PBST, for 1 hr, at room temperature. After washing twice in PBST, films were incubated with primary antibodies, overnight at 4 °C, in 3 wt% BSA/PBST. Following primary incubation and two PBST washes, secondary antibodies in PBS were incubated with the samples for 1–2 h at room temperature. Finally, samples were washed twice with PBS and left in a solution of PBS and DAPI stain (1:000 in PBS, ThermoFisher Scientific, 62248). Primary antibodies, and the dilutions used, were: β-tubulin (1:1500, TUJ1, Promega, G7121), Sox10 (1:300, Abcam, ab155279), and GFAP (1:500, DAKO, Z033429; EMD Millipore, AB5541). Secondary antibodies: Alexa Fluor 488 (1:1000, A11070), Alexa Fluor 647 (1:1000. A21236), Cy3-AffiniPure Donkey Anti-Chicken IgY (1:500, Jackson ImmunoResearch, 703-165-155).

Stained films were inverted onto a coverslip in a drop of Slow Fade Diamond Antifade Mountant (Invitrogen, S36972). Imaging was performed using a Leica DMi8 microscope, using an LASX software interface. With the high surface roughness of the films, z-stacks were taken at a minimum of 3 places on the film and post-processed with Thunder image analysis (Leica) to remove background fluorescence.

Scanning electron microscopy micrographs were taken of films without cells and after 1 week of co-culture. Samples with cells were dried in serial ethanol dilutions, at least 1 hour incubation in each: 70% ethanol, 80% ethanol, 90% ethanol, 100% ethanol. Finally samples were incubated in hexamethyldi-silazane (Sigma) for 10 minutes and allowed to air dry. Dried samples were adhered to 13mm aluminum stubs and sputter coated with platinum. Surfaces were examined using a Zeiss Auriga, at 7 keV in scanning electron mode, and 20 keV for backscatter electron images.

### 2.8 Micro Computed Tomography (μCT)

All tomography images were obtained using a Perkin-Elmer Quantum GX. Films were immersed in PBS, and imaged at 90 keV, 88 μA, with a 36 mm field of view at a 50μm resolution. After acquisition, x-ray attenuation was quantified in Image J. Values of attenuation were normalized by the attenuation of the native polymer film (0wt% TaO_x_), and are a composite measurement of the radiopaque matrix and the buffer present within the microporous regions of the film. Intensity was measured on 2-3 films from independent batches and are presented as mean ± standard deviation.

### 2.9 Statistics

All statistics were done using GraphPad Prism 9.4.1. Data was first analyzed via ANOVA, followed by multiple comparisons of the means, using Fisher’s LSD test. In all cases, α < 0.05 was considered significant, with a 95% confidence interval.

## 3 Results

### 3.1 Radiopaque films

Prior to testing biological response to radiopaque polymer substrates, films incorporating TaO_x_ nanoparticles were produced, and materials and imaging properties were characterized, Figure 1 (a). While the nominal incorporation was 0-40 wt% TaO_x_, when assessed via thermogravimetric analysis (TGA), the true range of TaO_x_ was between 0 - 36wt% TaO_x_. A minimum of 5wt% TaO_x_ was required to radiologically discriminate the films from buffer within a hydrated environment, simulating normal physiological conditions (phosphate buffered saline (PBS)). As the weight percentage of TaO_x_ nanoparticles increased, a corresponding increase in x-ray attenuation was observed when imaged using μCT, Figure 1(a-b). The distribution of nanoparticles was uniform throughout the films, imparting homogeneous x-ray attenuation to films. The measured attenuation was a composite value of both the radiopaque polymer matrix and the buffer trapped within the film microporous structure, which was normalized to have an attenuation of 0 HU. Overall, for each percent of TaO_x_ added, the attenuation of the films in buffer increased by 100 HU. At the nanometer scale, with nanoparticle addition, surface roughness was also significantly increased. Prior to nanoparticle addition, all polymer film surfaces were smooth, Figure 1(c1-e1), but reached a maximum surface roughness at 40wt% TaO_x_, Figure 1(c2-e2). Regardless of surface roughness, glia could interact and adhere to the film surface, Figure 2(e).

**Figure 1:**
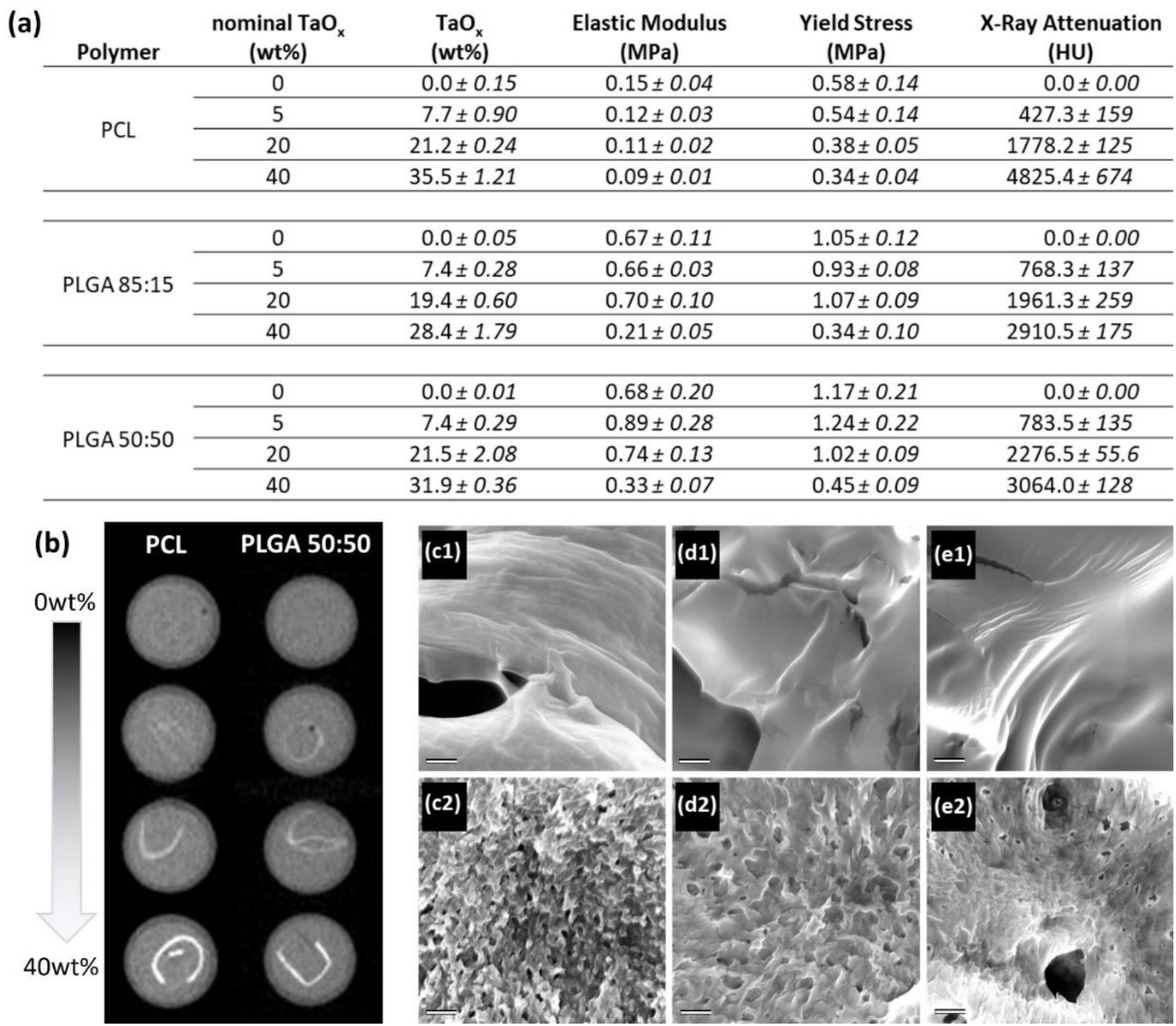
Porous polymer films were created incorporating 0-40wt% radiopaque nanoparticles, which (a) altered hydrated film properties including mechanics and x-ray attenuation. (b) X-ray attenuation was increased with increased TaO_x_ in the matrices, as shown via representative samples of PCL and PLGA 50:50; from top to bottom: 0, 5, 20, 40wt% TaO_x_. (c) Nanoparticles also corresponded to nano-scale changes of the surface as seen via scanning electron microscopy of (c) PCL (d) PLGA 50:50 and (e) PLGA 85:15 with (1) 0wt% TaO_x_ and (d) 40wt% TaO_x_. Scale bar (c-e): 500nm.

**Figure 2:**
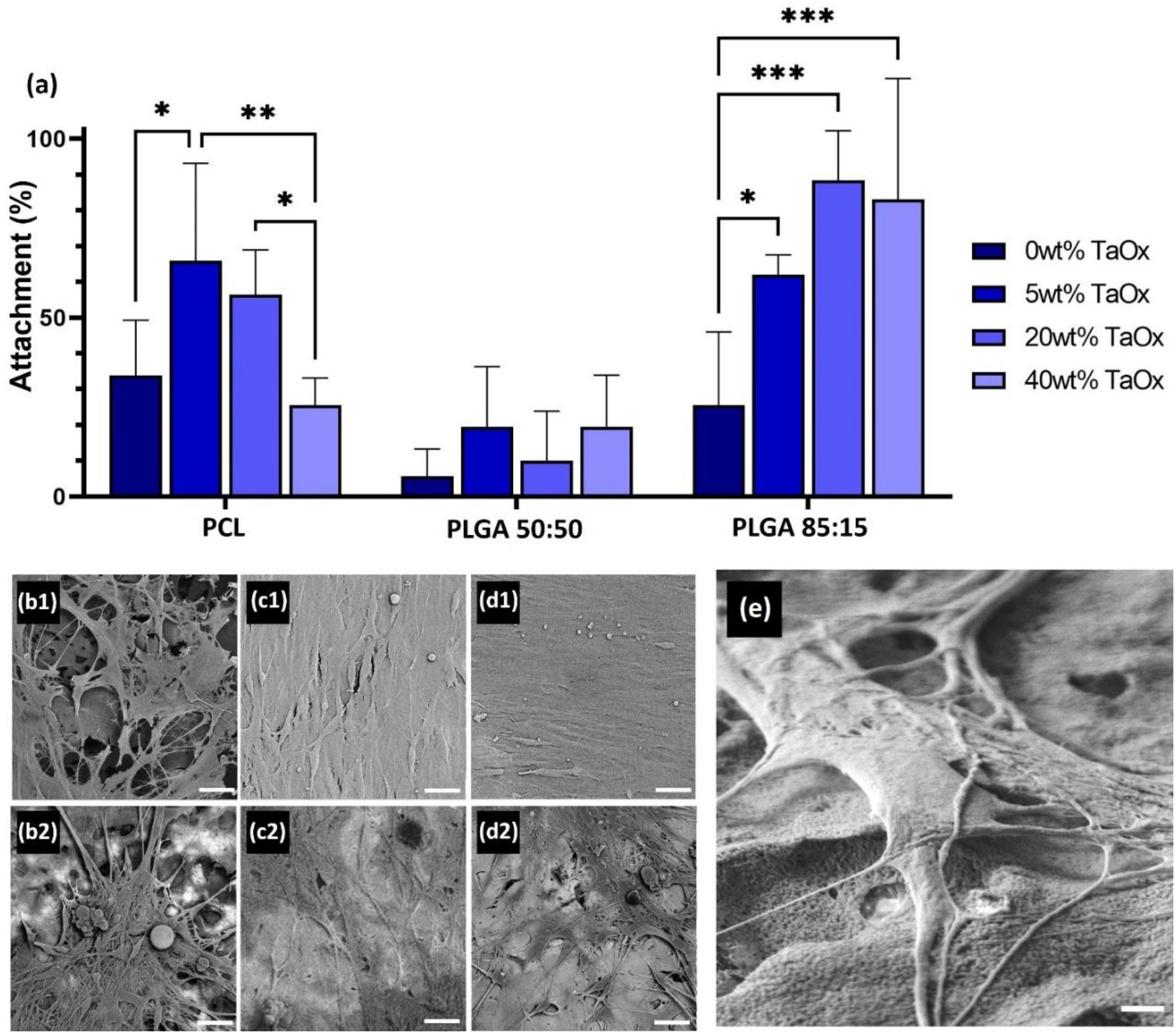
Glial cells attached and proliferated on all radiopaque films. (a) Attachment was affected by the type of polymer matrix and the weight percentage of TaO_x_. After one week, a co-culture of glia and DRG neurons proliferated, as seen via scanning electron microscopy of (b) PCL, (c) PLGA 50:50 and (d) PLGA 85:15 incorporating (1) 0wt% TaO_x_ and (2) 40wt% TaO_x_. (b-d) Micrographs were collected in backscatter mode, with lighter areas of the film having higher TaO_x_ content. Glia did not preferentially adhere or avoid regions of higher TaO_x_ in the films, but interacted directly with the film surface, as visualized on (e) PCL + 40wt% TaO_x_, imaged using secondary electron mode at high resolution. Scale bar: (b-d): 20 μm and (e) 2 μm. * < 0.05. ** < 0.01, *** < 0.001

At the macro-scale, mechanical testing of hydrated films also revealed a significant effect of TaO_x_ addition. With higher TaO_x_, the apparent elastic modulus and yield stress decreased significantly for all matrices. The trend was particularly noticeable in PLGA films, where both metrics decreased by over half between 20wt% and 40wt% TaO_x_. As expected, film porosity was the controlling factor of the apparent elastic modulus, with values of polymer without nanoparticles reaching only 0.15±0.04, 0.67±0.11 and 0.68±0.2 MPa for PCL, PLGA 85:15 and PLGA 50:50, respectively. PCL films had a significantly greater percentage elongation prior to failure compared to PLGA films, regardless of TaO_x_ addition. Percent elongation for PCL with 0wt% TaO_x_ was between 60-100% elongation and deceased to < 40% with 40wt% TaO_x_. PLGA 85:15 had the lowest percentage elongation (< 10%) and PLGA 50:50 reached up to 20-30% elongation, both of which decreased with increased TaO_x_.

### 3.2 Glial attachment

As the key cellular component orchestrating peripheral nerve repair, the effect of TaO_x_ incorporation on glial cell attachment was examined. After 24 hrs, even without TaO_x_ present, attachment on the polymer matrices was significantly different, with the lowest attachment on PLGA 50:50 (5.62 ±7.7%), Figure 2(a). In addition, TaO_x_ incorporation significantly affected attachment, although the trends observed were dictated by the polymer matrix, Figure 2. Porous PCL films showed an increase in attachment up to 5wt% TaO_x_, with a maximum at 65.8 ±27%, which then dropped significantly between 20 and 40wt% TaO_x_. In contrast, attachment on films of PLGA 85:15 increased significantly at all levels of TaO_x_ addition, while PLGA 50:50 films were unaffected by TaO_x_ nanoparticles.

Regardless of initial glial attachment, co-cultures of glia and DRG neurons were able to adhere and proliferate on all films. After one week of co-culture, a confluent cell layer was observed on all films with 0wt% TaO_x_, Figure 2(b1-d1). With increasing TaO_x_ content, cell confluency decreased on the film surfaces, with gaps between the cells devoid of glial or neurons, Figure 2(b2-d2). While the films had a homogeneous distribution of TaO_x_ nanoparticles, microscopy of the surface revealed that in PLGA films, the nanoparticles were more dispersed at the micro-scale. Despite PCL films having more concentrated areas of nanoparticles present, cells did not appear to preferentially adhere or avoid to the regions that were rich in TaO_x_ within the films.

### 3.3 Myelination potential on radiopaque films

For peripheral nerve repair, devices must promote myelination, the critical step in the repair process which leads to functioning nerves. Many of the key steps in myelination occur within glial cells, and are dependent on a highly coordinated cascade of transcription factors and specialized protein markers. The markers tested in this co-culture model of nerve repair were early markers of glial lineage (sox10) [18], transcription factors tied to initiating a “repair” phenotype (cJun) [19] and pro-myelination (Oct6) [20], as well as late markers of myelination (MPZ/P0) [21]. On all films tested, the myelination response of glia was affected by TaO_x_ incorporation, with significant changes occurring even with as little as 5wt% TaO_x_ incorporation. However, as with glial attachment, the three different polymer matrices showed different responses to nanoparticle addition.

When cultured on PCL films, co-cultures showed expression of all markers of myelination, Figure 3(a-b). The addition of 5wt% TaO_x_ appeared to slightly enhance the expression of these proteins, but this effect disappeared at > 20wt% TaO_x_. At 40wt% TaO_x_, nearly all myelination markers were down-regulated on the films. Markers expressed early in the repair process (sox10, c-Jun) were most significantly impacted. The expression data was consistent with changes in the co-culture on the PCL films, Figure 3(c-f). The confluent layer of elongated glia and neurites became clusters of isolated cells at 40wt% TaO_x_.

**Figure 3:**
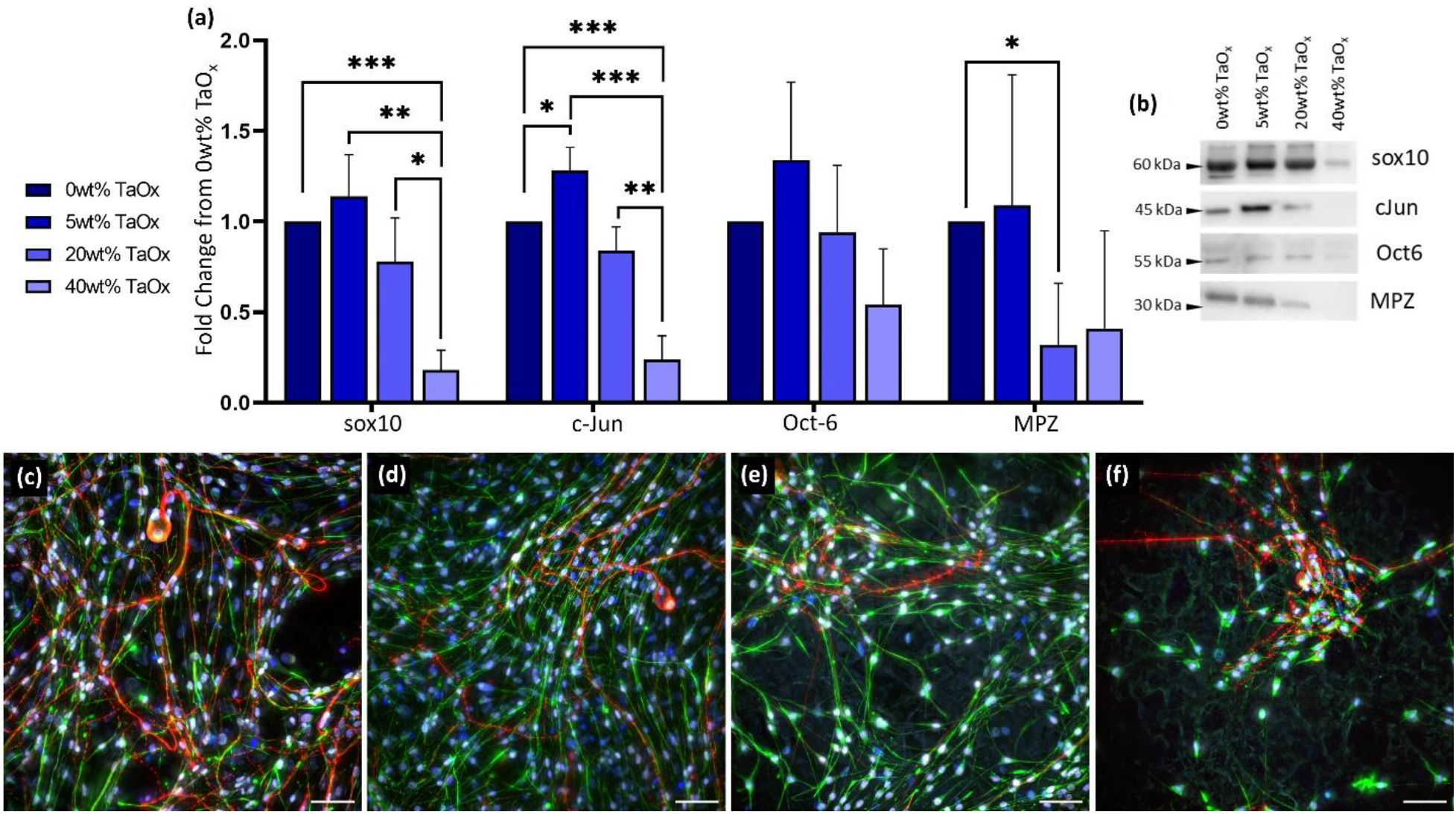
Glial proliferation and repair was affected by nanoparticle incorporation into a PCL matrix. (a) Expression of myelination markers were significantly affected with increasing TaO_x_ incorporation, as quantified by western blotting, with (b) showing typical expression of proteins as the weight percent of TaO_x_ changed. All expression data was normalized to 0wt% TaO_x_ films. (c-f) Fluorescent microscopy showed elongated glial cells and DRG neurites on PCL film surfaces with (c) 0wt% TaO_x_, (d) 5wt% TaO_x_, (e) 20wt% TaO_x_ and (f) 40wt% TaO_x_; red: DRGs (TUJ1), green: Schwanns (GFAP), white: sox10, blue: nuclei (DAPI). Scale bar (c-d): 50 μm. * p < 0.05. ** p < 0.01, *** p < 0.001

Markers of myelination were also present on PLGA films of different compositions. Co-cultures of glia and neurites created confluent layers of cell on both PLGA 85:15 (Figure 4(c1-f1)) and PLGA 50:50 (Figure 4(c2-f2)). However, expression of myelination markers differed between the two PLGA compositions. PLGA 85:15 had a trend for increased cJun and MPZ expression with 5-20wt% TaO_x_, despite no significant differences in the expression of sox10, Figure 4(a). In general, for PLGA 85:15, there was little or no difference between expression of markers on 0wt% and 40wt% TaO_x_. The high variability of the PLGA 85:15 samples precluded significant trends being made. Compared to native PLGA 50:50 films with 0wt% TaO_x_, co-cultures on radiopaque PLGA 50:50 substrates had a significantly decreased expression of sox10 and c-Jun, which remained consistent despite the weight percentage of TaO_x_ nanoparticles, Figure 4(b).

**Figure 4:**
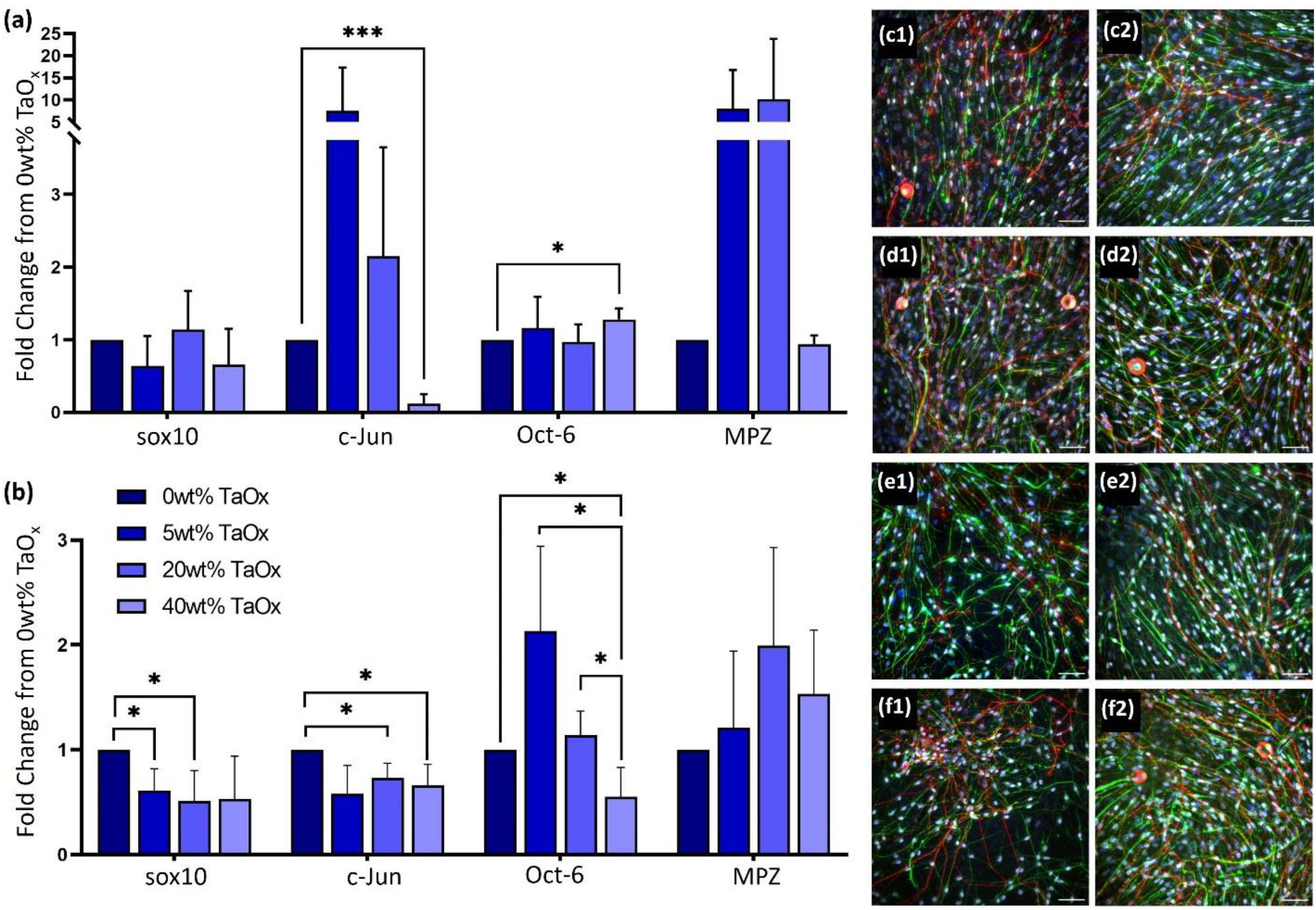
Myelination response of glial cells on PLGA films was dependent on TaO_x_ incorporation. Expression of myelination markers in the co-culture was investigated after one week on (a) PLGA 85:15 films and (b) PLGA 50:50 films containing 0-40wt% TaO_x_. All expression data was normalized to films with 0wt% TaO_x_. (c-f) Fluorescent microscopy showed elongated glial cells and DRG neurites on film surfaces with (c) 0wt% TaO_x_, (d) 5wt% TaO_x_, (e) 20wt% TaO_x_ and (f) 40wt% TaO_x_ in both (1) PLGA 85:15 and (2) PLGA 50:50; red: DRGs (TUJ1), green: Schwanns (GFAP), white: sox10, blue: nuclei (DAPI). Scale bar (c-f): 50 μm. * p < 0.05. ** p < 0.01, *** p < 0.001

Expression of Oct6 was also down-regulated at > 20wt% TaO_x_, but MPZ expression was unchanged by the addition of nanoparticles, even up to 40wt% TaO_x_.

Following the significant differences in glia attachment on the different polymer matrices, the expression of myelination markers was significantly different between radiopaque matrices, Figure 5. Overall, the marker for glial lineage, sox10 (Figure 5(a)) was lower on PLGA films than PCL films. In contrast, the late-stage myelin protein, MPZ, had significantly higher expression on PLGA 50:50 than all other matrices as the TaO_x_ incorporation was ≥ 20wt%, Figure 5(b). Expression reached 6.36±3.6 and 7.7±5.1 fold higher than PCL films in co-cultures on PLGA 50:50 with 20wt% and 40wt% TaO_x_, respectively. The expression pattern of myelination markers does not correlate with the mechanical properties of the films, Figure 1(a). Thus, the mode of cellular attachment to the film surfaces was investigated via the expression of integrin components α6 and β1, as integrin α6β1 is a laminin binding integrin that is highly expressed in glial cells [22]. When no nanoparticles were present in films (0wt% TaO_x_), expression of integrin α6 was down-regulated on PLGA 50:50, Figure 5(d). With 20wt% TaO_x_, there was a general down-regulation of integrin β1 on all matrices, although integrin α6 did not significantly change.

**Figure 5:**
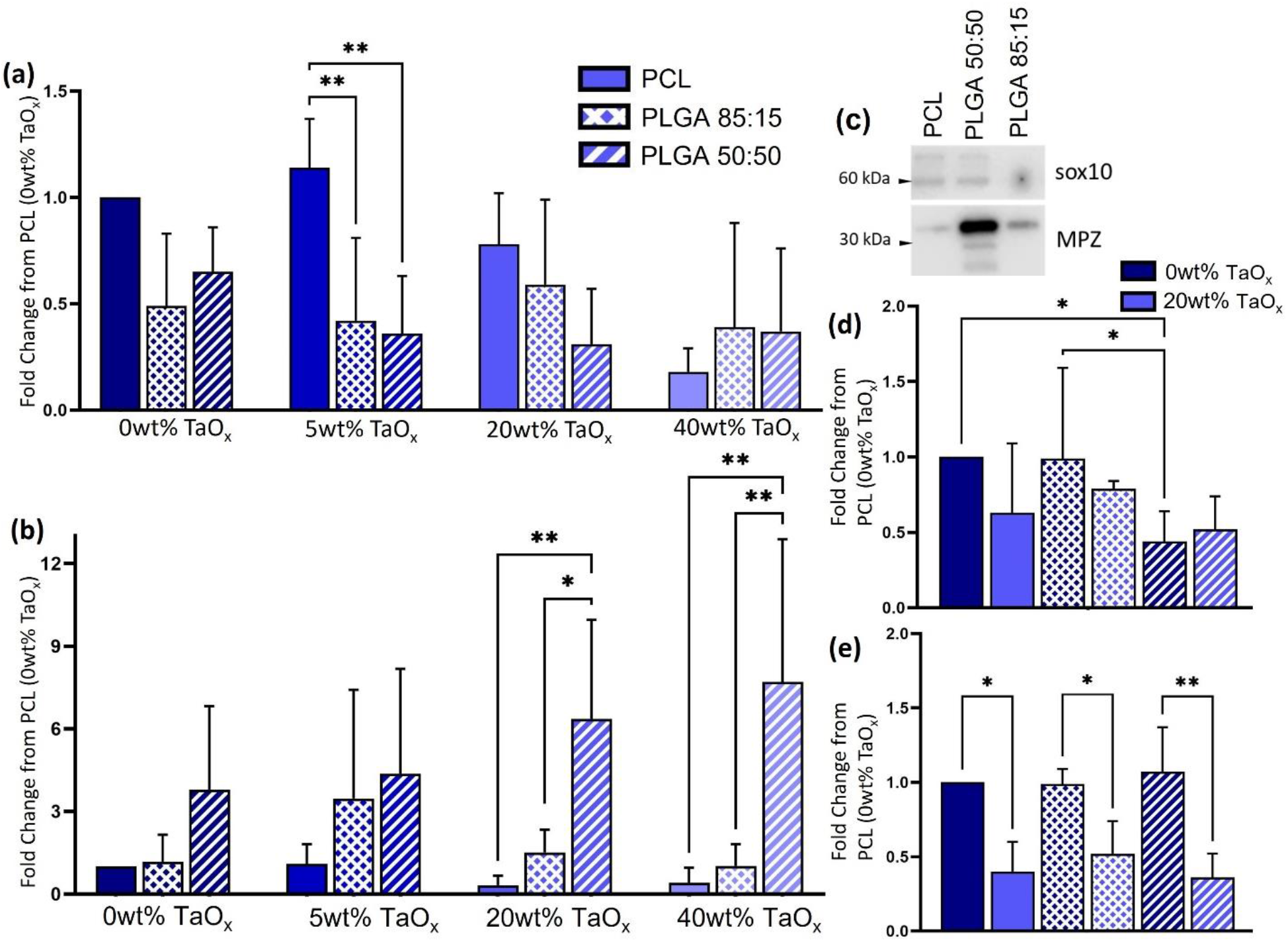
Myelination markers were significantly affected by the polymer matrix, with the effect of TaO_x_ incorporation dependent on the marker examined. (a) Glial lineage marker sox10 was generally down regulated on PLGA films, compared to PCL films. (b) In contrast, MPZ was up-regulated on PLGA 50:50, particularly at ≥ 20wt% TaO_x_, as (c) typical protein bands demonstrate on films with 40wt% TaO_x_. All protein expression data was reported as the fold change from expression on films of PCL + 0wt% TaO_x_. Cell attachment via integrin can mediate biological response. Expression of laminin integrins (d) α6 and (e) β1 were dependent on the polymer matrix and amount of TaO_x_ incorporated (0 vs 20wt% TaO_x_). *p < 0.05. ** p< 0.01

## 4 Discussion

The treatment of peripheral nerve injury remains an important clinical problem. Lack of autologous nerve tissue for autografts and concerns over immunological responses when using allograft tissue have driven research into alternative nerve repair devices, namely nerve conduits [14]. Conduits have been produced from many biocompatible, FDA approved polymers, including collagen, PCL and PLGA, and shown favorable clinical outcomes [5,14]. As the number of implanted biomedical devices used in the clinic grows, there is concern regarding the potential risks of failure after implantation. In situ radiological monitoring of devices is a promising way to assess long-term device function and enable prevention of catastrophic device failure in the clinic. Unfortunately, devices made from polymers are not radiologically distinguishable from surrounding tissues, requiring the introduction of radiopaque contrast agents, such as nanoparticles, to polymer devices [10,23,24]. However, with the creation of composite matrices of polymers and nanoparticles for peripheral nerve repair, it is important to balance the properties of the device that drive nerve repair and the introduction of radiopacity to enable longitudinal monitoring.

Tantalum based nanoparticles induce very low inflammatory responses and have demonstrated biocompatibility when used as an injected nanoparticle contrast agent [13,25]. Unlike injected nanoparticles which circulate freely and are quickly cleared from tissues and fluids, nanoparticles incorporated into biomedical devices are likely to be present at the target site for weeks to months and local concentration of nanoparticles will be affected by the degradation rate of the polymer. As the TaO_x_ nanoparticles are incorporated into a polymer matrix, they are shielded from direct contact with the cells. Thus, Schwann cell interactions with the radiopaque films are likely mediated via the polymer matrix properties and despite large changes in film properties, even with 40wt% TaO_x_ glial cells were able to attach and proliferate on all radiopaque polymer films.

### 4.1 Nerve repair on films

After traumatic injury of the peripheral nervous system, repair takes place in a coordinated, multi-stage process starting with the removal of damaged regions of nerve and the formation of a bridge of glial cells allowing regenerating neurites to span the damaged section [26]. The action of Schwann cells in the peripheral nervous system is central to recreating functioning nerve. During this process, Schwann cells must alter their gene expression, assuming a repair phenotype that is distinct from pro-myelinating immature Schwann cells [19,27]. Throughout development, myelination and repair, Schwann cells constitutively express sox10 (SRY-box transcription factor 10), a powerful transcription factor determining Schwann cell identity [28]. During the initiation of myelination in development, sox10 activates other key transcription factors, like Oct6, to turn on the expression of myelin proteins, such as myelin protein zero (MPZ) and myelin basic protein (MPB) [20]. In adult Schwann cells responding to injury, not only is Oct6 up-regulated, but the transcription factor c-Jun is also significantly up-regulated, and is believed to be critical for driving Schwann cell differentiation towards a repair phenotype [19]. As myelination progresses, Schwann cells wrap around neurites multiple times, nurturing the axon and insulating the electrical stimulation and creating a functional nerve [21].

To study the effect of biomedical devices on the repair process requires the utilization of a co-culture system of glia and neurons. Importantly, when attempting to draw conclusions about nerve repair in the clinic, the use of adult cells is critical. Co-culture models of myelination have been developed previously, showing myelination of adult neurons *in vitro* on porous polymer films mimicking biomedical devices [15]. This co-culture system relies on a layer of Schwann cells seeded onto the substrate mimicking the device prior to neurons. Unlike the previously reported model, which used a neonatal rat glia as the base layer, in the present case, all cells originated from post-natal mice [15]. Despite the adult origins of all cells, the mouse co-culture system showed universal expression of sox10 for glial lineage, early transcription factors for a repair phenotype (Oct6, c-Jun) and late markers of myelin (MPZ), Figures 3 and 4. When observed via microscopy, the glia were elongated and interacting closely with the neurites on all types of films, with and without radiopaque nanoparticles, suggesting that long-term monitoring of the peripheral nerve devices is feasible. However, the specific response of glial cells to TaO_x_ incorporation was dependent on the materials properties of the matrix polymer.

### 4.2 Properties of radiopaque films

The introduction of homogeneous radiopacity to synthetic polymer films was accomplished via biocompatible TaO_x_ nanoparticles with a hydrophobic coating. X-ray attenuation of the films was dependent on the weight percentage of TaO_x_ present and required a minimum of 5wt% to radiographically distinguish the films from their environment. The nominal amount of TaO_x_ was capped at 40wt% to retain as many properties of the polymer matrix as possible. The difference between the nominal and measured TaO_x_ at >20wt%, is likely due to loss of nanoparticles during the washing step, where sucrose was removed from the film to create the porous structure. At ≤ 20wt% TaO_x_, the difference in nanoparticle addition was within 10% between all polymer matrices.

To mimic implantable devices already in clinical use for the repair of peripheral nerves, films incorporated micro-scale porosity to facilitate nutrient diffusion, while maintaining the mechanical strength and compliance necessary to resist collapse [29,30]. However, mechanics and surface roughness are known modulators of cell behavior, and both metrics were affected by high weight percentages (> 20wt%) of TaO_x_ nanoparticle incorporation. While porosity is essential for nutrient diffusion through devices [30], the porosity introduced a micron scale surface roughness to the films [31]. The micron roughness was further modified with the development of nanoscale topography, particularly at 40wt% TaO_x_. While representing a subtle change in morphology, nanoscale roughness affects neural adhesion and growth [32], likely through protein adsorption [33].

Overall, the mechanical properties of the radiopaque films were determined by the polymer matrix, with PCL films exhibiting greater compliance than PLGA. However, with increasing TaO_x_ content, all films had significantly reduced mechanical properties and a lower percentage elongation, Figure 1(a). Even at low weight percentages of nanoparticle addition, there was no strengthening of the films, as seen in other nanoparticle composite systems [34,35]. The effect of nanoparticles on composite matrices is attributed to the interactions occurring at the interface of the polymer and nanoparticle. Where the interactions are weak, or agglomeration of the nanoparticles occurs, the overall mechanics of the composite are reduced [36]. In the case of PCL films, at high magnification, films with 40wt% TaO_x_ had regions of more concentrated nanoparticles visible at the surface, on the order of 10-40 μm; the size of these regions was small enough to appear homogeneous by CT. PLGA films in contrast were completely homogeneous, even at high resolution. Despite this, weak interactions at the interface may have allowed cavitation around the particles, leading to lower elastic modulus and an increasingly brittle behavior at high nanoparticle content.

### 4.3 Radiopacity supports myelination

As a hydrophobic, non-degrading synthetic polymer, PCL has a long history of biomedical use, including clinical nerve repair devices [4]. With the addition of TaO_x_ into PCL, the expression of both c-Jun and Oct6 decreased. Taken together with the lower expression of sox10 with increasing nanoparticle amount, the number and viability of glial cells was negatively affected by > 20wt% TaO_x_. Given the stability of PCL, which requires years to degrade in vivo [37], and the relative stability of the mechanical properties until 40wt% TaO_x_ addition, the gradual decrease in protein expression markers was likely due to the formation of nanoscale roughness on the film surface with TaO_x_ incorporation. Other studies have also observed decreased neuron and Schwann cell growth with nanoscale topography, with cells unaffected by microscale features [16,32].

Glial response to TaO_x_ nanoparticles was also dependent on the polymer matrix. On PLGA films, regardless of composition, sox10 expression was consistently down-regulated compared to PCL. This might indicate that a greater percentage of the cells present on the PCL films after one week co-culture were glial cells, as opposed to other proliferating cells like fibroblasts. While markers for glial lineage were down regulated, MPZ expression, indicating the formation of myelin, was promoted on PLGA 50:50 compared to all other matrices, particularly ≥ 20wt% TaO_x_. Up-regulation of MPZ would suggest that the glial cells present on the films are in a more advanced state of differentiation than those on PCL. A key difference between PCL and PLGA is in their mechanical properties. In the peripheral nervous system, stiffer matrices tend to push Schwann differentiation towards a myelinating phenotype in the presence of extra-cellular matrix cues [38]. However, all polymer matrices tested in the current study had an elastic modulus at least a hundred times higher than measured values of peripheral nerve tissue (50kPa), which in itself is not enough to drive Schwann differentiation [17], and differences in mechanical properties vanished at 40wt% TaO_x_ but not change in glial response. Aside from mechanics, degradation rates are also significantly different between PLGA and PCL. Overall, PLGA degrades faster, with the composition of PLGA giving insight into how degradation and degradation products may affect nerve repair.

Compositions of PLGA blends differ in their ratio of poly(lactic acid) to poly(glycolic acid). While PLGA composition did not affect the distribution of TaO_x_ nanoparticles in the film, or significantly impact the trends in surface roughness or mechanical properties, composition does affect hydrophobicity and degradation kinetics of matrices [39]. PLGA 85:15 degrades over several months, while PLGA 50:50, with the higher glycolic acid content, completely degrades within a few weeks. An accelerated degradation profile may indicate that glia have more exposure to both the TaO_x_ nanoparticles and the lactic acid and glycolic acid degradation products produced by PLGA. Pathways by which TaO_x_ interacts with cells have not been studied closely, but it is known that Schwann cell lactate metabolism, a pathway by which PLGA degradation products are consumed, is critical to the proper function of nerves [40]. Regardless, the effect of PLGA composition was apparent even at the earliest time points. Glial attachment, for example, was significantly lower on PLGA 50:50 than PLGA 85:15. Varied glial response continued after week 1, with decreased expression of sox10 and cJun in radiopaque PLGA 50:50 films balanced by up-regulation of Oct6 until > 20wt% TaO_x_ incorporation. MPZ expression was not significantly affected by TaO_x_ at all on PLGA 50:50, suggesting that the peak for the transiently expressed transcription factor c-Jun occurred earlier on films with increasing TaO_x_, to maintain levels of MPZ protein. Across the panel of expression markers, radiopaque PLGA 85:15 films showed the greatest variability in expression. With a decreased degradation rate compared to PLGA 50:50, other factors like surface roughness and mechanics likely played a more important and potentially conflicting role in biological response. There appeared to be a trend towards increased repair processes and myelination (cJun and MPZ) with 5-20wt% TaO_x_, but overall, there was little or no difference between expression of markers on 0 or 40wt% TaO_x_.

The specificity of myelination responses to the polymer films, indicate that different driving forces predominate in different polymer matrices. Key drivers of glial behavior include surface roughness, mechanics, surface chemistry and degradation products. In this study, polymer film surfaces were coated with laminin prior to co-culture, as the laminin has been shown to encourage neuronal outgrowth [15]. However, laminin also modulates Schwann cell behavior to environmental cues, such as mechanics [17,38]. Although not strictly necessary to achieve myelination, cellular pathways like YAP/TAZ transcription that mediate mechanosensing act via adhesion proteins, namely integrins [38]. Nanoscale surface roughness also acts on attachment by disrupting the formation of focal adhesion complexes through topographical cues or changes in the conformation of the proteins that adsorb to the surface [16,32]. Given the high likelihood that materials properties were being mediated by attachment, expression of integrin α6β1, the predominant attachment molecule to laminin in Schwann cells, was investigated [22].

The importance of laminin binding to radiopaque surfaces was seen in the deceased expression of integrin β1 as TaO_x_ incorporation increased from 0 to 20wt%. Blocking integrin β1 down-regulates Schwann cell attachment [22]. It is known that all three polymers have differences in hydrophobicity and degradation profiles and products. The down regulation of integrin β1 with higher TaO_x_ content was universal across the polymer matrices, making it likely that this was related to increases in surface roughness, a trait shared between all film types. Given the co-culture system, the expression of laminin integrins may be due to more than just the glial response alone, but we expect that Schwann cells will be the predominant cell type.

Integrin expression, while important, is likely not the only cellular pathway affected, as integrin expression alone cannot account for all changes in glial expression. While surface roughness likely played a large role on attachment, and therefore expression for the radiopaque films, differences in the expression of glial integrins occurred between PLGA 50:50 and PCL matrices even without nanoparticles present, shown as a significant down regulation of integrin α6. Surface chemistry, such as end groups of polymers, are also known to affect factors like cellular motility via integrin and focal adhesion kinases [41]. Over and above the surface chemistry, PLGA exhibits a fast degradation profile with the potential release of degradation products that could contribute to Schwann and neurite metabolism. The consistent up-regulation of MPZ on PLGA 50:50 films is most likely linked to these factors rather than integrin-mediated factors, as the level of both integrins on PLGA 50:50 and PCL was not significantly different at 20wt% TaO_x_, but expression of MPZ was.

In order to image peripheral nerve repair devices via CT, the process of nerve repair must be able to tolerate the addition of at least 5wt% TaO_x_ into the device matrix. For all matrices tested, the repair process was unimpeded up to 20wt% TaO_x_, satisfying this criterion. Regarding polymer choice for peripheral nerve repair, the complex interplay between different matrix signals demonstrates the importance of choosing an appropriate biomaterial for specific applications. This study raises the question of whether the apparent shift towards high expression of myelin proteins on PLGA 50:50 was due to a greater exposure of glia to TaO_x_ or to degradation products. In either case, there may be further therapeutic effects of radiopaque films to investigate in the future.

## 5 Conclusions

Incorporating radiopacity into implanted peripheral nerve conduits allows for long-term radiological monitoring, which has clinical value to diagnose and predict implant failure and enable earlier intervention before catastrophic consequences. With the addition of TaO_x_ nanoparticles, radiopacity was introduced to FDA-approved polymers, already used for nerve repair conduits. As the amount of TaO_x_ reached 40wt%, surface roughness of the films increased at the nanoscale, along with a significant decrease in mechanical strength. Radiopaque films were able to support the attachment and myelination response of Schwann cells, controlled primarily by different properties of the film matrices. An optimum range of 5-20wt% TaO_x_ was found to allow for films to be radiographically distinguished from background tissue, but still encourage the maximum myelination response. This defines a usable range of radiopaque TaO_x_ nanoparticles, for use in nerve repair devices and validates the use of long-term monitoring for this application.

## 6 Acknowledgments

The authors would like to thank Foster Buchanan for supplying the nanoparticles and Per Askeland from the MSU Composite Materials and Structures Center for help with scanning electron microscopy. This study was supported by the National Institute of Biomedical Imaging and Bioengineering of the NIH under award number R01EB029418. The content is solely the responsibility of the authors and does not necessarily represent the official views of the National Institutes of Health.

## Data Availability

Data is available on request to the authors.

