## Supplemental for "Radiopaque Implantable Biomaterials for Nerve Repair"

Western blotting was used to quantify protein expression. Co-cultures were lysed in Ripa buffer (supplemented with phosphatase inhibitors and protease inhibitors), and spun down at 15,000×g for 5 min at 4 °C. After which, supernatants were collected and the protein concentration of the supernatant was measured (BioRad, BCA kit). Supernatants were combined with 2× Laemmli buffer, incubated at 60°F for 20 min, separated by SDS/ PAGE (4 µg of protein per lane), and transferred to PVDF membranes. Transfer was accomplished using the iBlot 2 Gel Transfer Device (invitrogen, IB21001), a dry transfer system, at the recommended setting of 20 V for 1 min, 23 V for 4 min and 25 V for 2 minutes (7 min total). PVDF membranes were blocked with 5% dried milk in PBST (phosphate buffered saline pH 7.4, containing 0.1% Tween-20) and probed with primary antibodies specific for Oct-6 (1:1,000, Aviva Systems Bio, ARP33061_T100), MPZ (1:1000; Aves, PZO), c-Jun (1:1000, Cell Signaling Technology, 60A8), Sox10 (1:1000, Abcam, ab155279), integrin α6 (abcam, ab181551), and integrin β1 (abcam, ab179471) at 4 °C overnight. The next day, the membrane was washed and incubated with anti-chicken IgY-HRP (12-341, Millipore) or anti-rabbit IgG-HRP (12–348, Millipore, ab205718, abcam) at room temperature for 2 hr and further washed in PBST six times. Early problems with dark spots on developed membranes were traced to agglomeration of the secondary anti-rabbit IgG-HRP (12–348, Millipore) after storage; from that point, the anti-rabbit IgG-HRP ab205718 (abcam) was used exclusively.

Protein bands were visualized with ECL Plus substrate (cytiva, RPN2232) on a Li-Cor Odyssey FC and resulting signal was quantified via Image J. Membranes were then washed in PBS and stripped for 15 minutes in Restore Western Blot Stripping Buffer (Thermo Scientific 21059) with subsequent PBS washing. Membranes were either blocked and reprobed or were stained for total protein using the BLOT-Fast Stain (G Biosciences, cat# 786-34) according to manufacturer's directions. The total protein signal was imaged under visible light on a c300 imager (azure biosystems).

Signal for total protein in each lane was quantified on Image J, and the protein loading was normalized to the signal from the first lane. The normalized signal was used to adjust the measured band intensity from antibody staining, also quantified using Image J. For comparison of nanoparticle effect, all protein expression was recorded as a fold change from matrices with 0wt% $TaO_x$. The final protein quantification reported is the result of three biological replicates (2 technical replicates each). Samples of PCL + 0wt% $TaO_x$ were present on all blots and used to normalize differences between membrane exposure.

In the following figures, each set of membranes is presented with all protein bands shown, along with the total protein staining for each membrane. For integrin expression, only bands at high molecular weight, corresponding to the mature protein, were quantified. The molecular weight of bands on integrin blots was determined from an initial blot, with a molecular weight marker present.

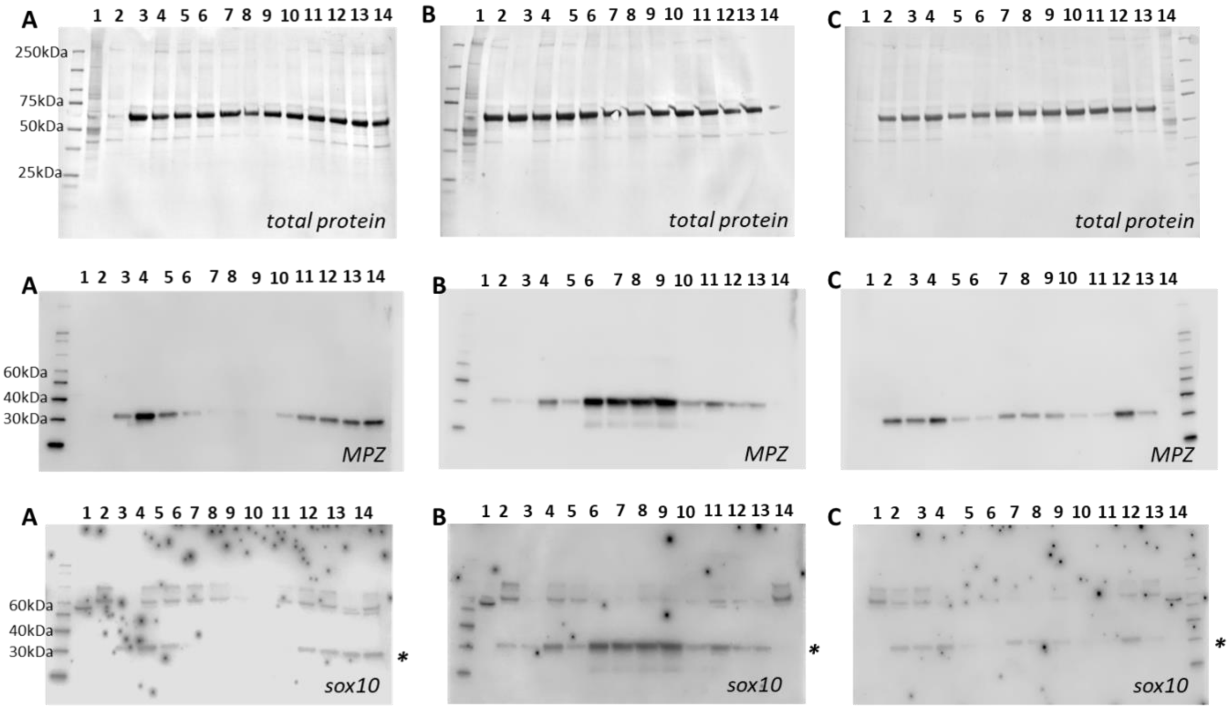


**A**

**1:** brain lysate
**2:** glass
**3:** PLGA 85:15 (0wt% $TaO_x$)
**4:** PLGA 50:50 (0wt% $TaO_x$)
**5:** PCL (0wt% $TaO_x$)
**6:** PCL (5wt% $TaO_x$)
**7:** PCL (20wt% $TaO_x$)
**8:** PCL (40wt% $TaO_x$)
**9:** PCL (40wt% $TaO_x$)
**10:** PCL (20wt% $TaO_x$)
**11:** PCL (5wt% $TaO_x$)
**12:** PCL (0wt% $TaO_x$)
**13:** PLGA 85:15 (0wt% $TaO_x$)
**14:** PLGA 50:50 (0wt% $TaO_x$)

**B**

**1:** brain lysate
**2:** PCL (0wt% $TaO_x$)
**3:** PLGA 85:15 (0wt% $TaO_x$)
**4:** PLGA 50:50 (0wt% $TaO_x$)
**5:** PLGA 50:50 (5wt% $TaO_x$)
**6:** PLGA 50:50 (20wt% $TaO_x$)
**7:** PLGA 50:50 (40wt% $TaO_x$)
**8:** PLGA 50:50 (40wt% $TaO_x$)
**9:** PLGA 50:50 (20wt% $TaO_x$)
**10:** PLGA 50:50 (5wt% $TaO_x$)
**11:** PLGA 50:50 (0wt% $TaO_x$)
**12:** PCL (0wt% $TaO_x$)
**13:** PLGA 85:15 (0wt% $TaO_x$)
**14:** glass

**C**

**1:** glass
**2:** PLGA 50:50 (0wt% $TaO_x$)
**3:** PCL(0wt% $TaO_x$)
**4:** PLGA 85:15 (0wt% $TaO_x$)
**5:** PLGA 85:15 (5wt% $TaO_x$)
**6:** PLGA 85:15 (20wt% $TaO_x$)
**7:** PLGA 85:15 (40wt% $TaO_x$)
**8:** PLGA 85:15 (40wt% $TaO_x$)
**9:** PLGA 85:15 (20wt% $TaO_x$)
**10:** PLGA 85:15 (5wt% $TaO_x$)
**11:** PLGA 85:15 (0wt% $TaO_x$)
**12:** PLGA 50:50 (0wt% $TaO_x$)
**13:** PCL (0wt% $TaO_x$)
**14:** brain lysate

Figure S1: Western blots of MPZ and sox10 expression (biological replicate 1). Top row: total protein, second row: MPZ staining, third row: sox10 staining, fourth row: sample labels. * Indicates a band remaining from previous staining, before stripping and reprobing the membranes.

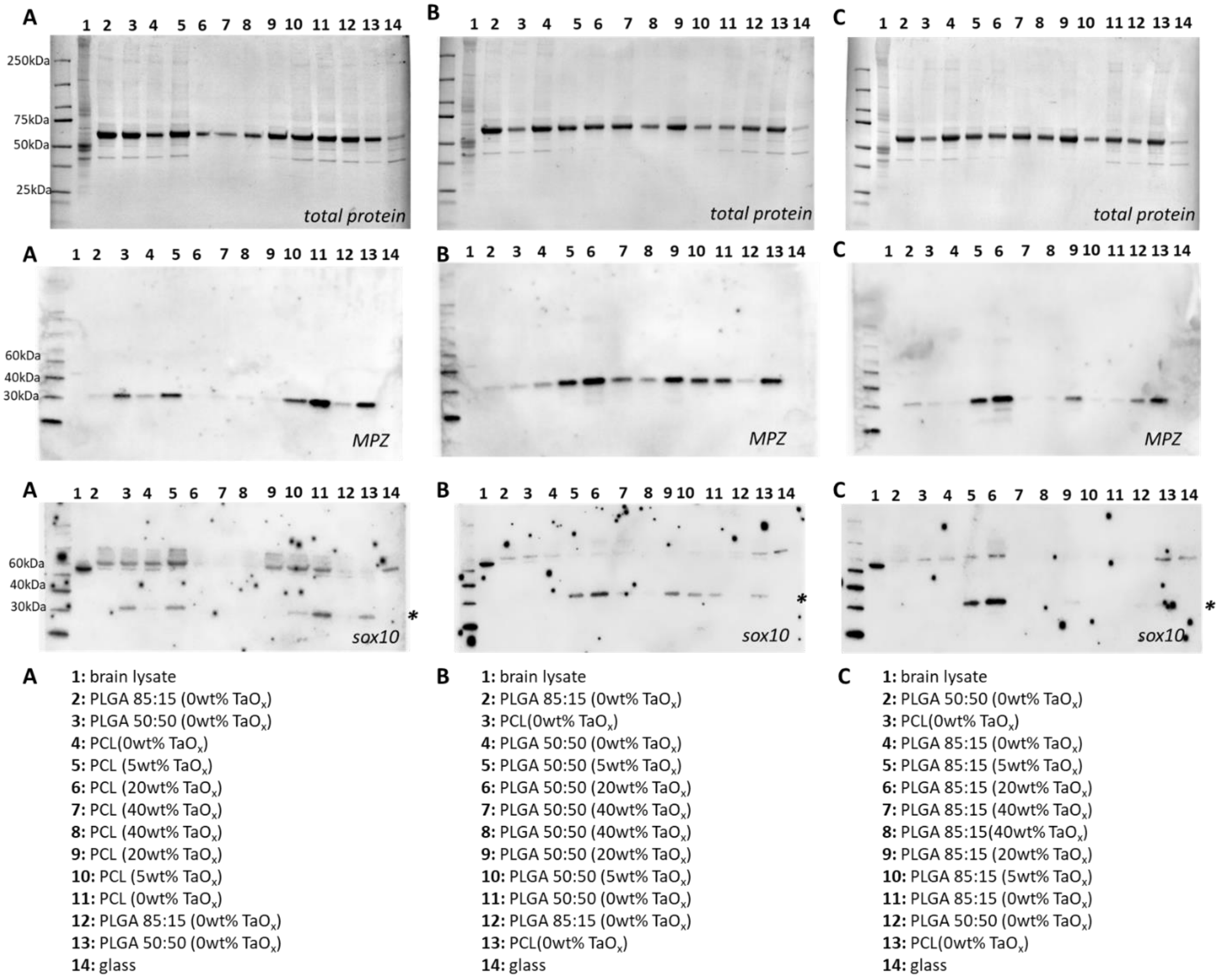


Figure S2: Western blots of MPZ and sox10 expression (biological replicate 2). Top row: total protein, second row: MPZ staining, third row: sox10 staining, fourth row: sample labels. * Indicates a band remaining from previous staining, before stripping and reprobing the membranes.

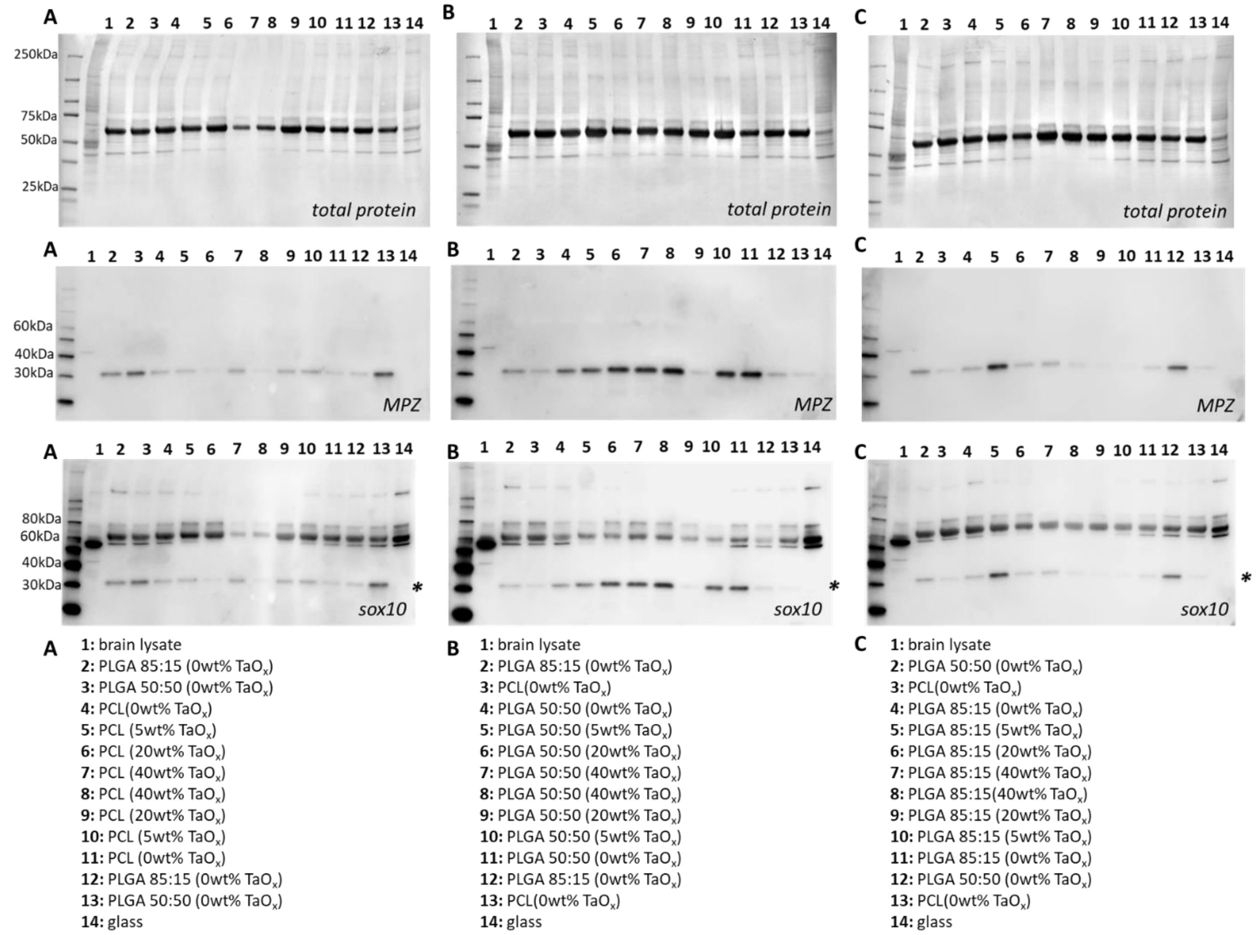


Figure S3: Western blots of MPZ and sox10 (biological replicate 3). Top row: total protein, second row: MPZ staining, third row: sox10 staining, fourth row: sample labels. * Indicates a band remaining from previous staining, before stripping and reprobing the membranes.

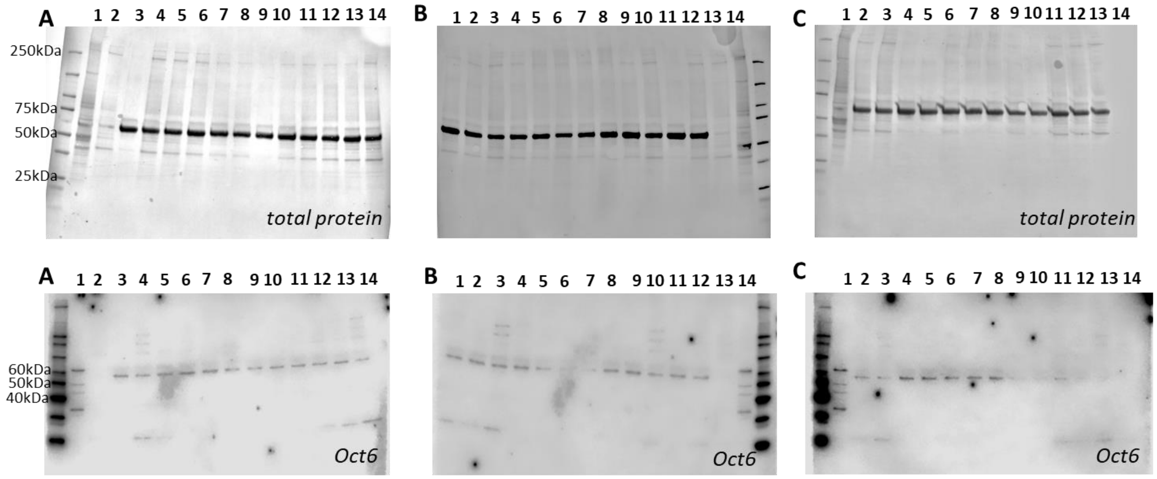


**A**
**1:** brain lysate
**2:** glass
**3:** PLGA 85:15 (0wt% $TaO_x$)
**4:** PLGA 50:50 (0wt% $TaO_x$)
**5:** PCL (0wt% $TaO_x$)
**6:** PCL (5wt% $TaO_x$)
**7:** PCL (20wt% $TaO_x$)
**8:** PCL (40wt% $TaO_x$)
**9:** PCL (40wt% $TaO_x$)
**10:** PCL (20wt% $TaO_x$)
**11:** PCL (5wt% $TaO_x$)
**12:** PCL (0wt% $TaO_x$)
**13:** PLGA 85:15 (0wt% $TaO_x$)
**14:** PLGA 50:50 (0wt% $TaO_x$)

**B**
**1:** PLGA 85:15 (0wt% $TaO_x$
**2:** PCL (0wt% $TaO_x$)
**3:** PLGA 50:50 (0wt% $TaO_x$)
**4:** PLGA 50:50 (5wt% $TaO_x$)
**5:** PLGA 50:50 (20wt% $TaO_x$)
**6:** PLGA 50:50 (40wt% $TaO_x$)
**7:** PLGA 50:50 (40wt% $TaO_x$)
**8:** PLGA 50:50 (20wt% $TaO_x$)
**9:** PLGA 50:50 (5wt% $TaO_x$)
**10:** PLGA 50:50 (0wt% $TaO_x$)
**11:** PLGA 85:15 (0wt% $TaO_x$)
**12:** PCL (0wt% $TaO_x$)
**13:** glass
**14:** brain lysate

**C**
**1:** brain lysate
**2:** PCL (0wt% $TaO_x$)
**3:** PLGA 50:50 (0wt% $TaO_x$)
**4:** PLGA 85:15 (0wt% $TaO_x$)
**5:** PLGA 85:15 (5wt% $TaO_x$)
**6:** PLGA 85:15 (20wt% $TaO_x$)
**7:** PLGA 85:15 (40wt% $TaO_x$)
**8:** PLGA 85:15(40wt% $TaO_x$)
**9:** PLGA 85:15 (20wt% $TaO_x$)
**10:** PLGA 85:15 (5wt% $TaO_x$)
**11:** PLGA 85:15 (0wt% $TaO_x$)
**12:** PCL (0wt% $TaO_x$)
**13:** PLGA 50:50 (0wt% $TaO_x$)
**14:** glass

Figure S4: Western blots of Oct6 expression (biological replicate 1). Top row: total protein, second row: Oct6, third row: sample labels.

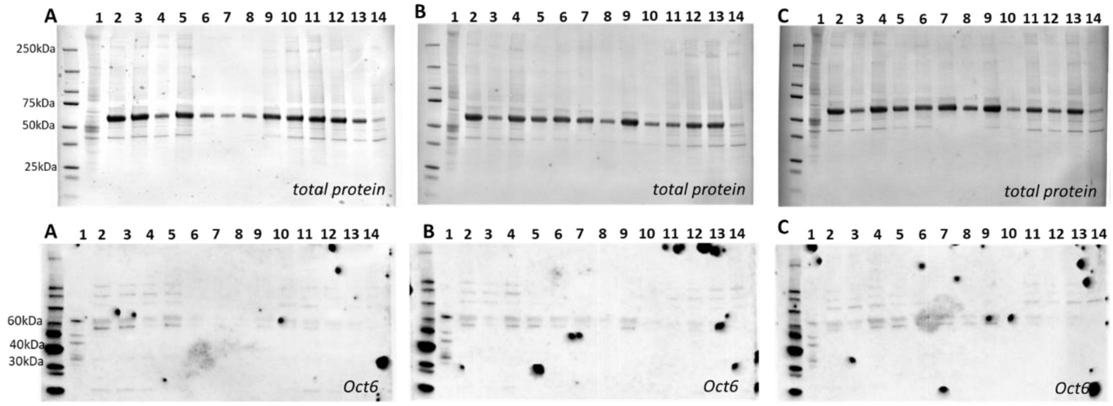


**A**
**1:** brain lysate
**2:** PLGA 85:15 (0wt% $TaO_x$)
**3:** PLGA 50:50 (0wt% $TaO_x$)
**4:** PCL(0wt% $TaO_x$)
**5:** PCL (5wt% $TaO_x$)
**6:** PCL (20wt% $TaO_x$)
**7:** PCL (40wt% $TaO_x$)
**8:** PCL (40wt% $TaO_x$)
**9:** PCL (20wt% $TaO_x$)
**10:** PCL (5wt% $TaO_x$)
**11:** PCL (0wt% $TaO_x$)
**12:** PLGA 85:15 (0wt% $TaO_x$)
**13:** PLGA 50:50 (0wt% $TaO_x$)
**14:** glass

**B**
**1:** brain lysate
**2:** PLGA 85:15 (0wt% $TaO_x$)
**3:** PCL(0wt% $TaO_x$)
**4:** PLGA 50:50 (0wt% $TaO_x$)
**5:** PLGA 50:50 (5wt% $TaO_x$)
**6:** PLGA 50:50 (20wt% $TaO_x$)
**7:** PLGA 50:50 (40wt% $TaO_x$)
**8:** PLGA 50:50 (40wt% $TaO_x$)
**9:** PLGA 50:50 (20wt% $TaO_x$)
**10:** PLGA 50:50 (5wt% $TaO_x$)
**11:** PLGA 50:50 (0wt% $TaO_x$)
**12:** PLGA 85:15 (0wt% $TaO_x$)
**13:** PCL(0wt% $TaO_x$)
**14:** glass

**C**
**1:** brain lysate
**2:** PLGA 50:50 (0wt% $TaO_x$)
**3:** PCL(0wt% $TaO_x$)
**4:** PLGA 85:15 (0wt% $TaO_x$)
**5:** PLGA 85:15 (5wt% $TaO_x$)
**6:** PLGA 85:15 (20wt% $TaO_x$)
**7:** PLGA 85:15 (40wt% $TaO_x$)
**8:** PLGA 85:15(40wt% $TaO_x$)
**9:** PLGA 85:15 (20wt% $TaO_x$)
**10:** PLGA 85:15 (5wt% $TaO_x$)
**11:** PLGA 85:15 (0wt% $TaO_x$)
**12:** PLGA 50:50 (0wt% $TaO_x$)
**13:** PCL(0wt% $TaO_x$)
**14:** glass

Figure S5: Western blots of Oct6 expression (biological replicate 2). Top row: total protein, second row: Oct6, third row: sample labels.

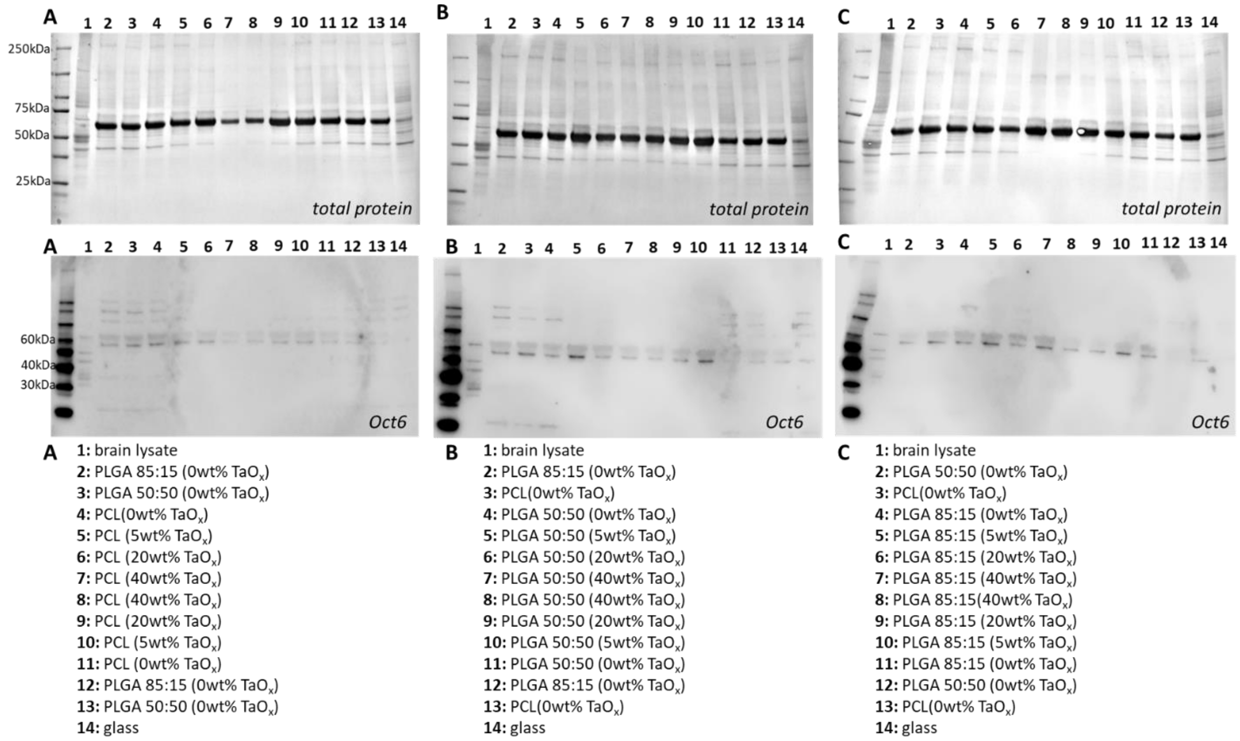


Figure S6: Western blots of Oct6 expression (biological replicate 3). Top row: total protein, second row: Oct6, third row: sample labels.

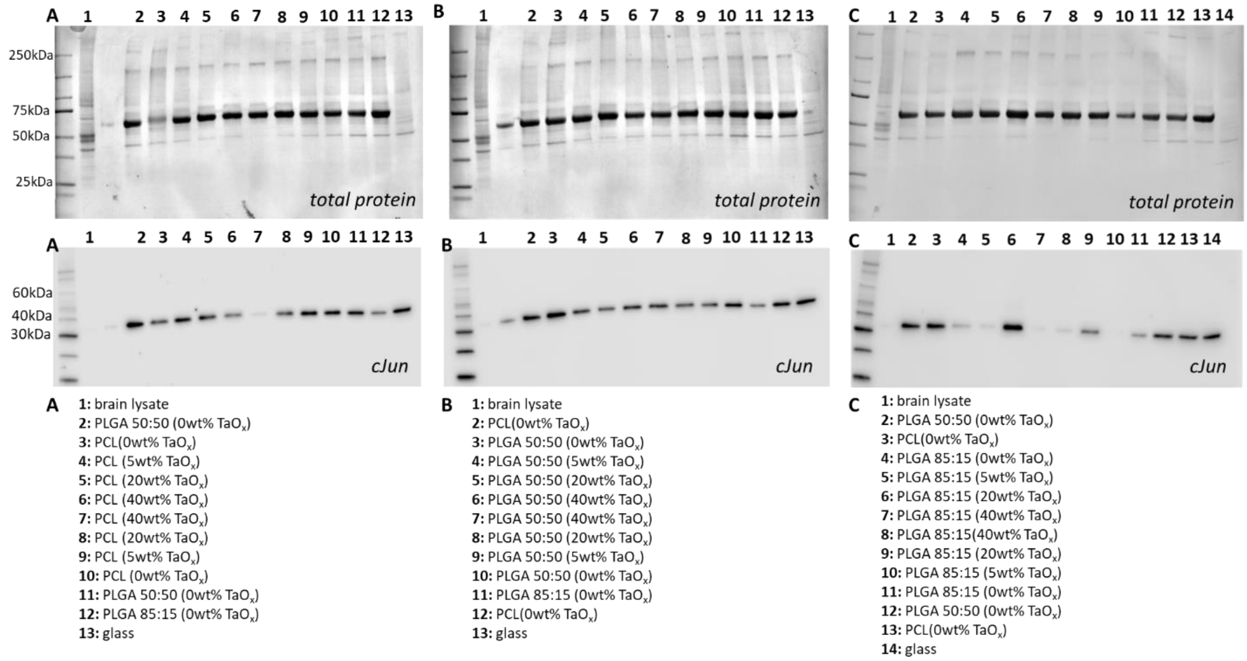


Figure S7: Western blots of cJun expression (biological replicate 1). Top row: total protein, second row: cJun, third row: sample labels.

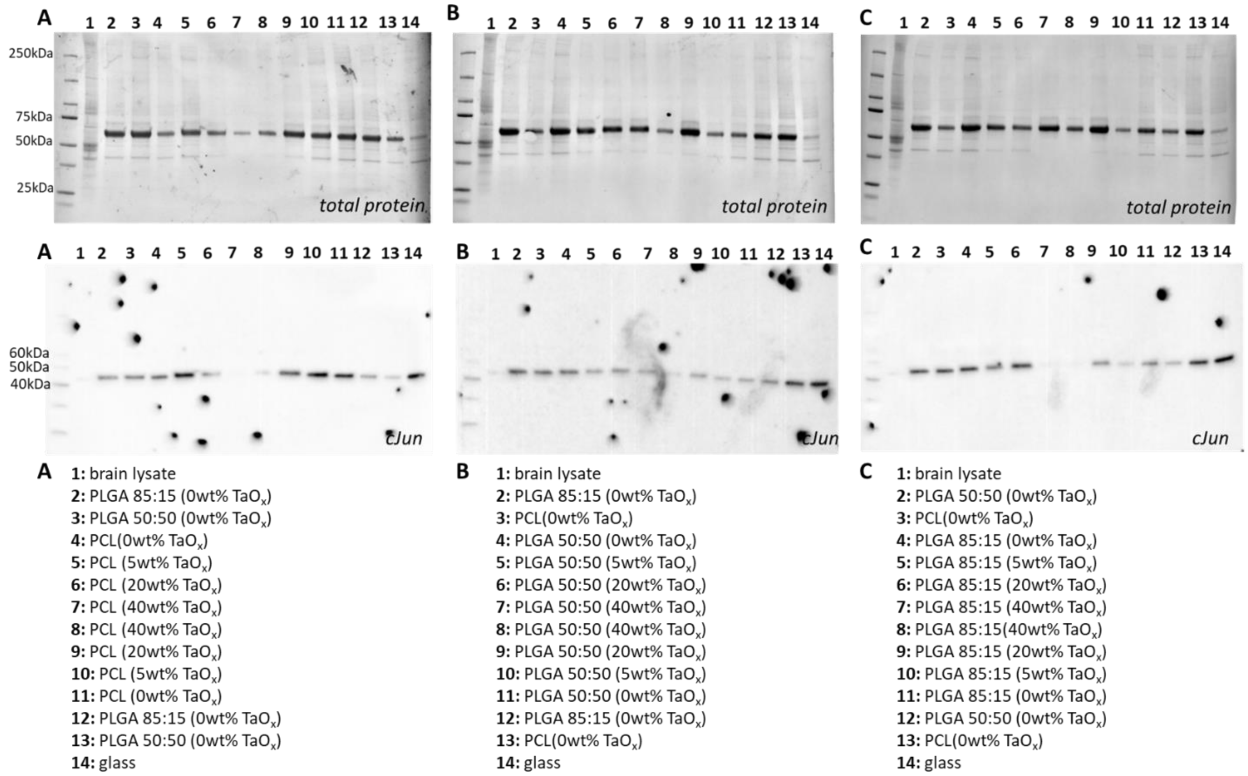


Figure S8:  Western blots of cJun expression (biological replicate 2). Top row: total protein, second row: cJun, third row: sample labels.

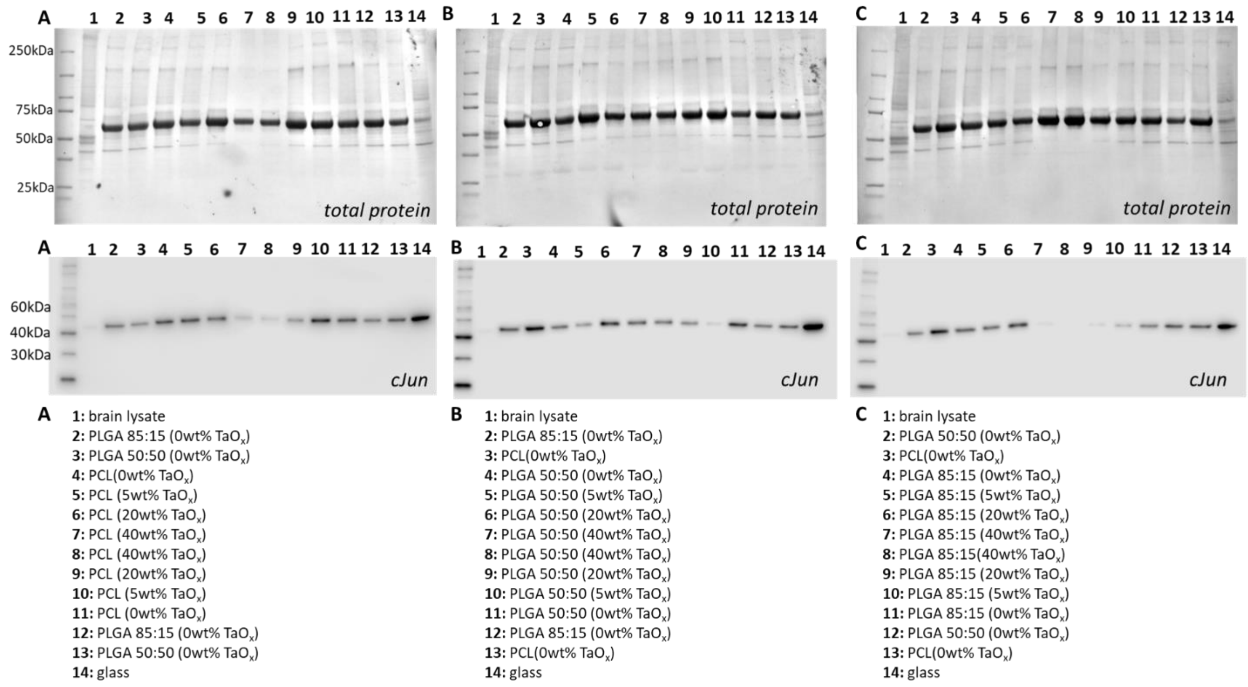


Figure S9:  Western blots of cJun expression (biological replicate 3). Top row: total protein, second row: cJun, third row: sample labels.

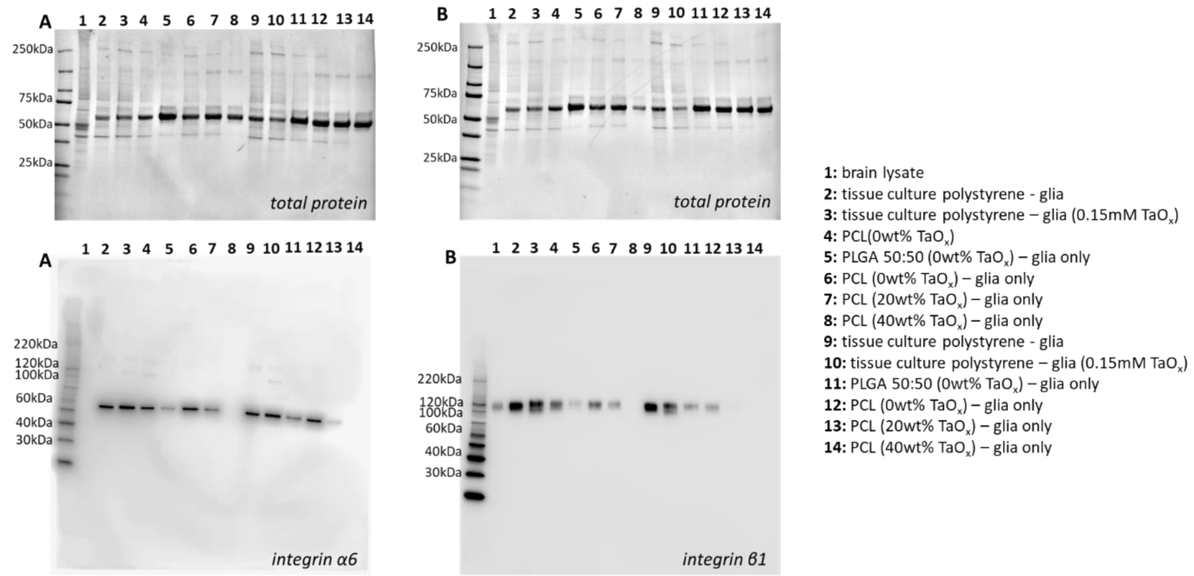


Figure S10: Western blots of integrin α6 and integrin β1 to establish whether glia had measurable expression. Top row: total protein, second row: (A) integrin α6 (B) integrin β1. Samples labels are the same for both membranes. Faint bands are visible at high molecular weight on integrin α6 membrane, corresponding to the full-length protein.

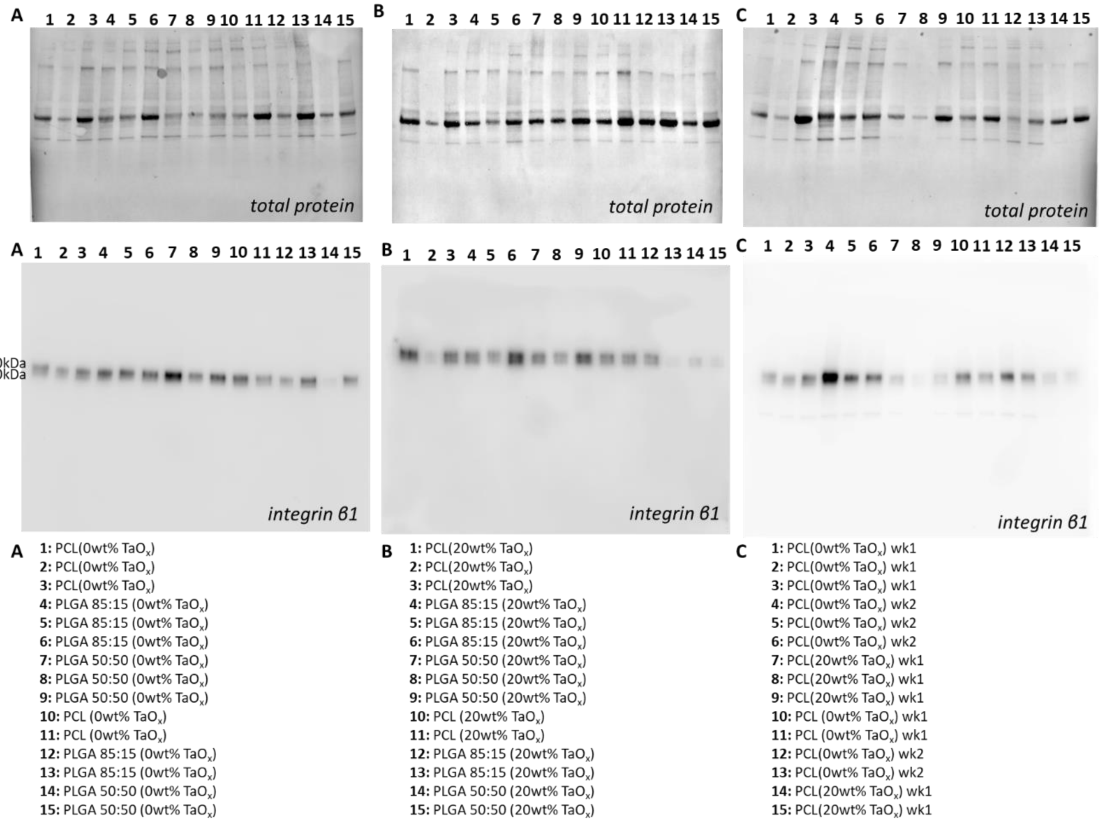


Figure S11: Western blots of integrin β1 expression in co-cultures of Schwann cells and neurons. Top row: total protein, second row: integrin β1 expression on films with (A) 0wt% $TaO_x$, (B) 20wt% $TaO_x$ and (C) a comparison between samples, third row: sample labels.

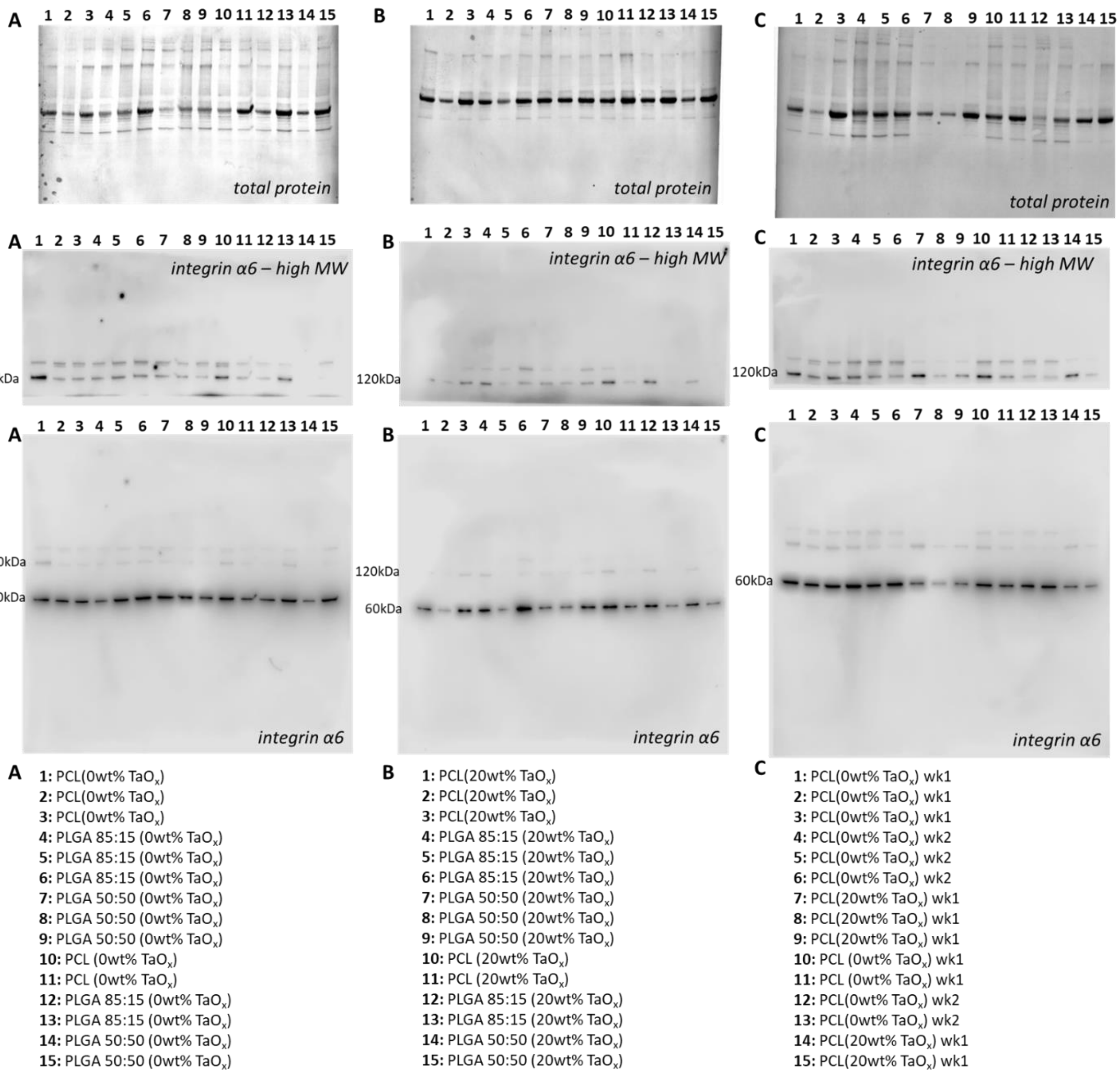


Figure S12: Western blots of integrin α6 expression in co-cultures of Schwann cells and neurons. Top row: total protein, second and third row: integrin α6 expression on films with (A) 0wt% $TaO_x$, (B) 20wt% $TaO_x$ and (C) a comparison between samples, fourth row: sample labels. Only high molecular weight bands (second row) were used for quantification, as these were associated with full length mature protein.
